# Cell wall pectic β-1,4-galactan contributes to increased plant freezing tolerance induced by cold acclimation

**DOI:** 10.1101/2023.05.31.542803

**Authors:** Daisuke Takahashi, Kouichi Soga, Kazuma Sasaki, Takuma Kikuchi, Tatsuya Kutsuno, Yui Nishiyama, Kim L. Johnson, Arun Sampathkumar, Antony Bacic, Toshihisa Kotake

**Author notes:** Daisuke Takahashi Graduate School of Science & Engineering, Saitama University, 255 Shimo-Okubo, Sakura-ku, Saitama 338-8570, Japan **Email:**. **Author Contributions:** Research ideas and design were completed by D. T. and T. Ko. Experiments were performed and data analysis was completed by D.T., K.So, K.Sa, T.Ki., T. Ku., and Y.N. Data interpretation and manuscript writing were completed by D.T., K.L.J., A.S., A.B. and T. Ko.

## Abstract

Subzero temperatures are often lethal to plants. Many temperate plants have a mechanism called cold acclimation that allows them to sense a drop in temperature and prepare for freezing stress through accumulation of soluble sugars and cryoprotective proteins. Plant cells are surrounded by a cell wall and as ice formation primarily occurs in the extracellular matrix (the apoplast), cell wall properties are important for plant freezing tolerance. Although previous studies have shown that the amounts of constituent sugars of the cell wall, in particular those present in pectic polysaccharides, are altered by cold acclimation, the significance of this change during cold acclimation has not been clarified. We found that β-1,4-galactan, which forms neutral side chains of pectin, accumulates in the cell walls of Arabidopsis and various freezing tolerant vegetables during cold acclimation. The *gals1 gals2 gals3* triple mutant, which has reduced β-1,4-galactan in the cell wall, exhibited reduced freezing tolerance compared with wild-type Arabidopsis during initial stages of cold acclimation. Expression of genes involved in the galactan biosynthesis pathway such as *GALACTAN SYNTHASES* and *UDP-glucose 4-epimerases* were induced during cold acclimation in Arabidopsis explaining galactan accumulation. Cold acclimation resulted in a decrease in extensibility and an increase in rigidity of the cell wall in the wild type, whereas these changes were not observed in the *gals1 gals2 gals3* triple mutant. These results indicate that the accumulation of pectic β-1,4-galactan contributes to acquired freezing tolerance by cold acclimation *via* changes in cell wall mechanical properties.

## Introduction

Plants experience environmental stress and it is freezing stress that most easily has lethal consequences. This is because cold temperatures not only reduce intracellular activity, but also lead to ice nucleation on the outside of cells that additionally exposes them to dehydration and mechanical stress (1, 2). One example of the tremendous impact of cold on plant ecosystems is the spring freeze that occurred in North America in 2007. This cold spell caused $2.7 billion in agricultural damage in the Central Plains and Midwest of the USA and a variety of other adverse ecological effects (3–6). Damage from freezing in Canada and other northern regions may increase as global warming progresses (7).

Because land plants, unlike animals, have limited capacity for movement or temperature regulation, and, unlike microorganisms, have complex multicellular tissues connected by cell walls, they are thought to need their own mechanisms for sensing temperature drops and controlling freezing injury. In fact, when exposed to non-freezing low temperatures, many plants are able to enhance freezing tolerance, a process referred to as cold acclimation (CA). During CA, a cellular protection mechanism is activated, which includes increased gene expression of transcription factors (e.g. *CBFs/DREBs*) and their downstream genes (e.g. *CORs*, *GolSs*, and *RD29s*), and accumulation of soluble sugars and membrane protective proteins (8). CA regulatory genes are being tested as candidate frost tolerance genes with the aim of breeding hardier barley (9).

In addition to these known mechanisms, recent studies have shown that cell wall mechanisms are necessary for enhancement of freezing tolerance during CA (10–17). The cell wall plays an important role in determining plant shape and mechanical properties during growth in plants exposed to a wide range of environments (18). In addition, the cell wall, which encases cells, also serves as a dynamic primary barrier that is important for their survival strategies. In freezing stress, ice nuclei formed in tissues are usually located in the apoplast (the cell wall space). Therefore, when cells are injured by freezing, the ice usually penetrates and/or strongly deforms the cell wall (11). Changes in the structure and properties of the cell wall are thus believed to be important in the control of tissue freezing (11, 13).

The primary cell wall in dicotyledonous plants is composed of pectin, hemicellulose, and cellulose together with small amounts of (glyco)proteins (18, 19). Cellulose is a main load-bearing structure in the cell wall, but other components also affect the mechanical properties of the cell wall. In walls of dicots, pectin is a key component of the matrix and influences both the mechanical properties and the porosity of the cell wall. Pectin is categorized into three main types, homogalacturonan (HG), rhamnogalacturonan (RG)-I, and RG-II (Fig. S1) (20). The HG region in pectin forms calcium cross-links, which strengthen the network structure of the cell wall. Calcium cross-linking of HG is believed to increase with cold acclimation and its importance in the acquisition of freezing tolerance has been demonstrated in various plants (15, 17, 21, 22). RG-II forms boron cross-links, and they have been demonstrated to be required for basal freezing tolerance before CA (10). The backbone of RG-I is composed of repeating units of D-galacturonic acid (GalA) and L-rhamnose (Rha) residues (23) and decorated with side chains, the two major types of which are neutral chains of α-1,5-arabinan and (arabino-)β-1,4-galactan (Fig. S1) composed of L-arabinose (Ara) and D-galactose (Gal), respectively. Arabinan has been implicated in the stomatal opening mechanism and in the regulation of physical properties of inflorescence stems through the control of cell wall flexibility (24–26). By contrast, the galactan biosynthetic mutant, *galactan synthase1* (*gals1*) *gals2 gals3*, exhibits little phenotypic difference to wild type (WT) under normal conditions and thus the physiological importance and function(s) of galactan are still elusive. Pectic galactan may be required under certain stressed conditions, including freezing (27–29).

Several studies have reported that Gal content is increased in pectin during CA (12, 14, 30–33). However, direct evidence regarding the accumulation of pectic galactan in CA, and its involvement in improving freezing tolerance is rudimentary. In this study the CA-induced increase in content of pectic galactan was thoroughly investigated. An increase in pectic galactan during CA was found in several freezing tolerant vegetables, including the model plant *Arabidopsis thaliana*. It contributed to plant freezing tolerance and was accompanied by changes in the mechanical properties of the cell wall. Our findings indicate that compositional modification of the cell wall during CA is important for improving freezing tolerance through changes in cell wall properties, adding to the increasing body of evidence pointing to the importance of the plant cell wall in responding to external environmental challenges to ensure plant survival.

## Results

### Cold induced increases in Gal residues in the pectin component of cell walls are common in many temperate plants

We first examined pea (*Pisum sativum* ‘753’), spinach (*Spinacia oleracea* ‘Progress’), Japanese mustard spinach (*Brassica rapa* var. *Perviridis* ‘Hamatsuduki’) and crown daisy (*Glebionis coronaria* ‘Satoakira’) from different families for freezing tolerance and observed a reduction of freezing injury after CA induction in all plants. (Fig. 1A). The cell wall changes in these plants were analyzed, focusing on the pectin-rich fraction (Fig. S2). Monosaccharide composition of pectin fractions revealed that the proportion of Gal increased in spinach, Japanese mustard spinach and crown daisy during CA (Fig. 1*B*). Unlike other vegetables, pea contained abundant Ara in the pectin fraction and similar proportions of Gal in non-acclimated (NA) and CA plants (Fig. 1*B*). However, since the amount of total pectin content increased in CA even for pea, the amount of Gal per fresh weight tended to increase during CA (Figs. S2 and S3). Thus, the absolute amount of Gal in the pectin fraction increased in all plants investigated in this study.

**Figure 1.**
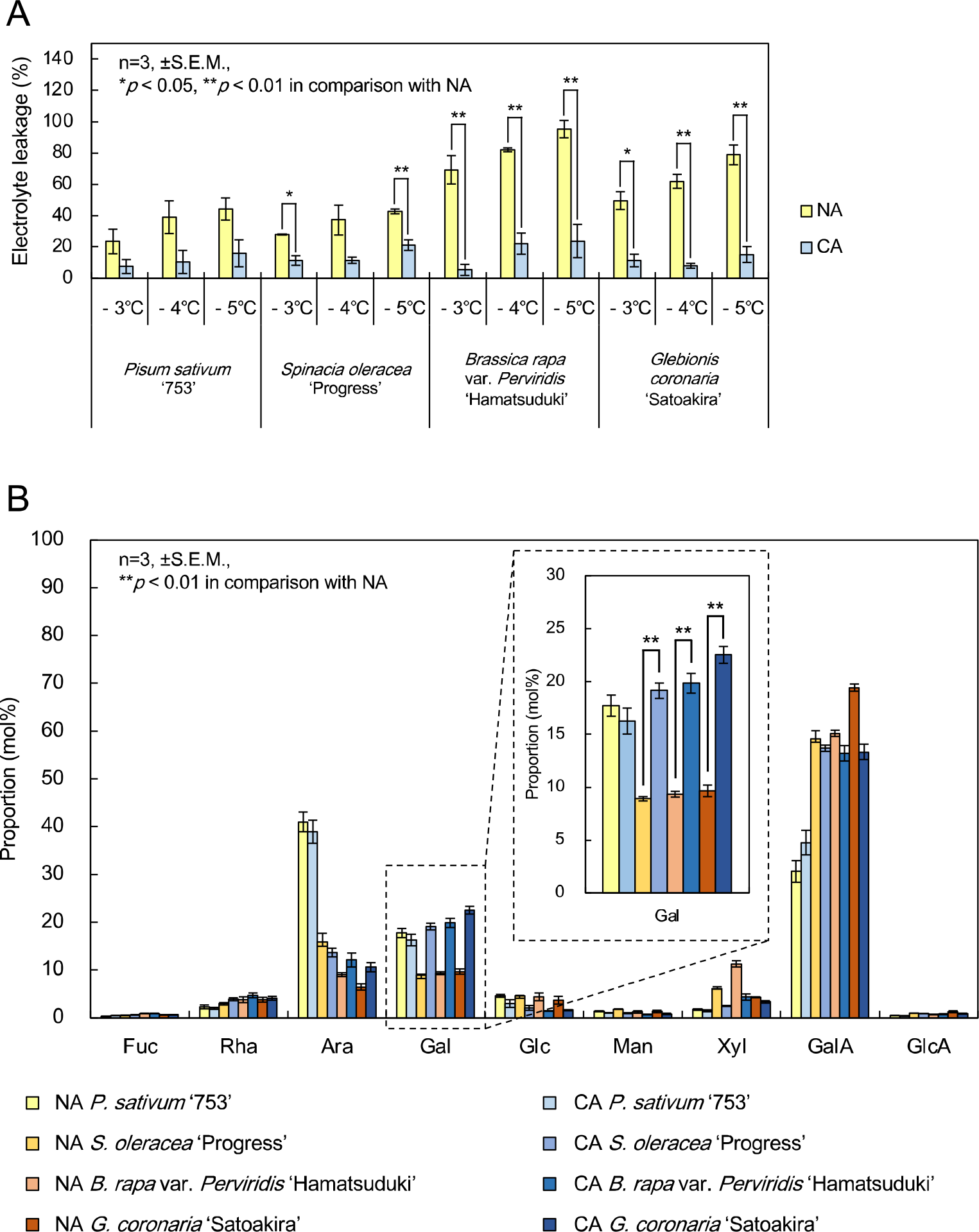
Plants with cold acclimation ability increase the galactose content of pectin under cold acclimation treatment. (A) Freezing tolerance of various vegetables in non-acclimation (NA) and cold acclimation (CA). After freezing at a certain freezing temperature, the electrical conductivity was measured before and after boiling, then the percentage of electrolyte leakage due to freezing injury was calculated. Statistically significant differences between NA and CA were determined with Student’s *t*-test (\**p* < 0.05, \*\**p* < 0.01). Error bars indicate ± S.E.M. (n = 3). (B) Monosaccharide composition in the pectin fraction of four vegetables before and after CA. The bar graph shows the percentage of each monosaccharide in pectin fractions. An enlarged view of galactose (Gal) is shown in the dotted box. (A and B) Statistically significant differences between NA and CA in Gal proportions were determined with Student’s *t*-test (\*\**p* < 0.01). Error bars indicate ± S.E.M. (n = 3). Fuc, fucose; Rha, rhamnose; Ara, arabinose; Gal, galactose; Glc, glucose; Man, mannose; Xyl, xylose; GalA, galacturonic acid; GlcA, glucuronic acid.

### Impaired β-1,4-galactan accumulation during CA in the *gals* triple mutant

Increased Gal content in the pectin fraction is presumed to be mostly due to accumulation of pectic galactan. Therefore, we analyzed pectic galactan accumulation during CA in the model plant *Arabidopsis thaliana* and its mutants *gals1 gals2 gals3* by several different methods. In Arabidopsis wild type (WT) plants, a significant increase in the proportion and content of Gal in the pectin-rich EDTA fraction occurred with CA as it did in the vegetables (Fig. 2*A* and *B* and Fig. S4). The increase of Gal in CA was suppressed in *gals1 gals2 gals3* compared to the WT (Fig. 2B). To confirm whether pectic galactan was indeed increased in WT, β-1,4-galactan in the pectin-rich EDTA fraction was specifically degraded with β-1,4-galactanase, and its hydrolysis product, galactobiose (Gal_2_), was quantified by HPLC (Fig. 2*C* and *D*). Pectic galactan increased over 7 d of CA in WT and was barely detectable in *gals1 gals2 gals3*. Furthermore, the relative amount of β-1,4-galactan in the pectin fraction was quantified by glycosidic linkage analysis (Fig. S5) and the data showed that indeed linear β-1,4-galactan was increased in CA and extremely low in the triple mutant. In addition, pectic galactan quantification in a series of homozygous single and double *gals* mutants was also performed. Pectic galactan levels were the same or higher in single mutants, and slightly less in both *gals1 gals2* and *gals2 gals3* double mutants compared to WT (Fig. S6). Therefore, no one specific GALS contributes to pectic galactan accumulation during CA, but rather the three GALSs encoded by the Arabidopsis genome seem to work in concert. Furthermore, observation of the tissue distribution of pectic galactan using the LM5 antibody, which specifically recognizes β-1,4-galactan (Fig. S1), showed that it accumulates throughout the leaf, including the epidermis and mesophyll cells, upon CA.

**Figure 2.**
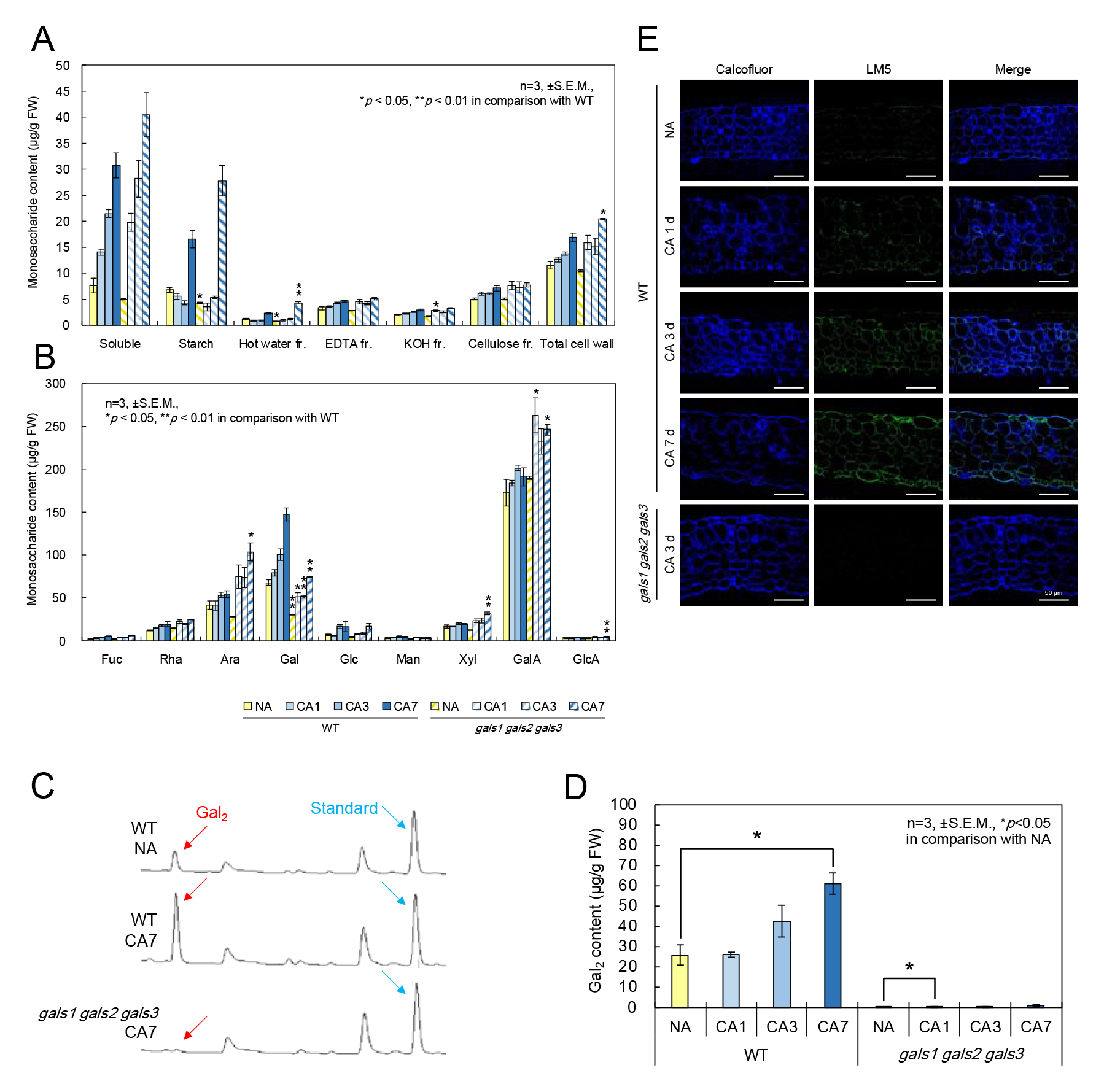
**Accumulation of pectic galactan in *Arabidopsis thaliana*** (A) Proportions of each fraction after the sequential extraction of the cell wall. Arabidopsis was ground and fractionated into a soluble sugar fraction, a starch fraction, hot water and EDTA fractions containing pectin, a KOH fraction containing hemicellulose, and a cellulose fraction containing crystalline cellulose. The respective sugar content per fresh weight is indicated. The rightmost bar graph shows the sum of the hot water, EDTA, KOH, and cellulose fractions containing cell wall components. (B) Monosaccharide composition of the pectin-rich EDTA fraction extracted by cell wall fractionation. (C) Pectin fractions extracted from Arabidopsis were subjected to β-1,4-galactanase, which specifically degrades pectic galactan, and the released galactobiose (Gal_2_) was detected by HPLC. Blue arrows indicate maltopentaose added as internal standard. (D) The amount of pectic galactan per fresh weight (FW) was estimated from the Gal_2_ peak area detected in C. (A, B and D) Values represent the amount of each sugar per FW. Statistically significant differences between NA and other treatments were determined with Dunnett’s test (\**p* < 0.05). Error bars indicate ± S.E.M. (n = 3). Fuc, fucose; Rha, rhamnose; Ara, arabinose; Gal, galactose; Glc, glucose; Man, mannose; Xyl, xylose; GalA, galacturonic acid; GlcA, glucuronic acid. (E) Immunohistochemical staining using antibody LM5, which specifically recognizes pectic galactan. Leaf sections of wild type (WT) and pectic galactan synthesis mutant (*gals1 gals2 gals3*) were prepared and tissue distribution of pectic galactan was observed. Blue indicates Calcofluor staining and green indicates LM5 staining. Scale bars, 50 µm.

### CA-induced freezing tolerance is reduced in an Arabidopsis the *gals* triple mutant

To confirm that pectic galactan does indeed affect freezing tolerance, we examined the freezing tolerance of *gals1 gals2 gals3* triple mutants with impaired pectic galactan synthesis in the model plant *Arabidopsis thaliana* (Fig. 3*A* and *B*). The survival curve obtained by the electrolyte leakage method showed no significant difference between WT and *gals1 gals2 gals3* in NA. However, at several temperature points in CA for 1 d (CA1) and 3 d (CA3), the percentage of electrolyte leakage was significantly higher in *gals1 gals2 gals3* than in WT. In CA for 7 d (CA7), *gals1 gals2 gals3* became as tolerant to freezing as the WT at all freezing temperatures. This indicates that acquisition of freezing tolerance is less likely to occur in the *gals1 gals2 gals3* compared to WT at the initial stage of CA. To verify whether freezing tolerance of *gals1 gals2 gals3* was reduced in specific organs, live tissues in aerial organs were stained red with 2,3,5-triphenyltetrazolium chloride (TTC) after freeze-thawing (Fig. 3*B*). No significant differences were observed between WT and *gals1 gals2 gals3* in NA, while a decrease in freezing tolerance was observed at CA3 compared to WT for the entire *gals1 gals2 gals3* plants. These data complement the immunohistochemistry data (Fig. 2*E*) thereby suggesting that pectic galactan contributes to CA-induced acquisition of freezing tolerance in whole leaves. To understand which of three galactan synthases, GALS1, GALS2, and/or GALS3 contributes to the increase in freezing tolerance by CA, we evaluated freezing tolerance using a series of single and double mutants (Fig. S7) and found that none of the single gene mutants were significantly different from WT in NA. After CA3, however, the *gals1 gals2 gals3* triple mutant alone was significantly less tolerant to freezing than WT indicating that all GALS enzymes contribute in a coordinated manner to increased freezing tolerance during the CA process.

### Various galactan-related genes are induced during the cold acclimation process

Galactan synthesis is achieved by GALS1, 2 and 3 which are glycosyltransferases catalyzing transfer of Gal residues from UDP-Gal in the Golgi apparatus (Fig. S1) (27, 28, 34) and UDP-glucose 4-epimerases (UGEs) which control the level of the nucleotide sugar donor UDP-Gal (Fig. 4*A*). UGEs are widely conserved in land plants, including monocots and mosses (Fig. S9) (35). Phylogenetic analysis of the GT92 family containing GALS homologous proteins in various land plants led to division into three clades, two consisting of dicotyledonous and one of monocotyledonous plants, and a loose grouping of other land plants (Fig. S8). In Arabidopsis, UGE2 is the major enzyme in forming UDP-Gal from UDP-Glc, whereas UGE1 and UGE3 found in other subclade are presumed to have different functions from UGE2 (36–38). UGE4 shares high sequence similarity with UGE2.

**Figure 3.**
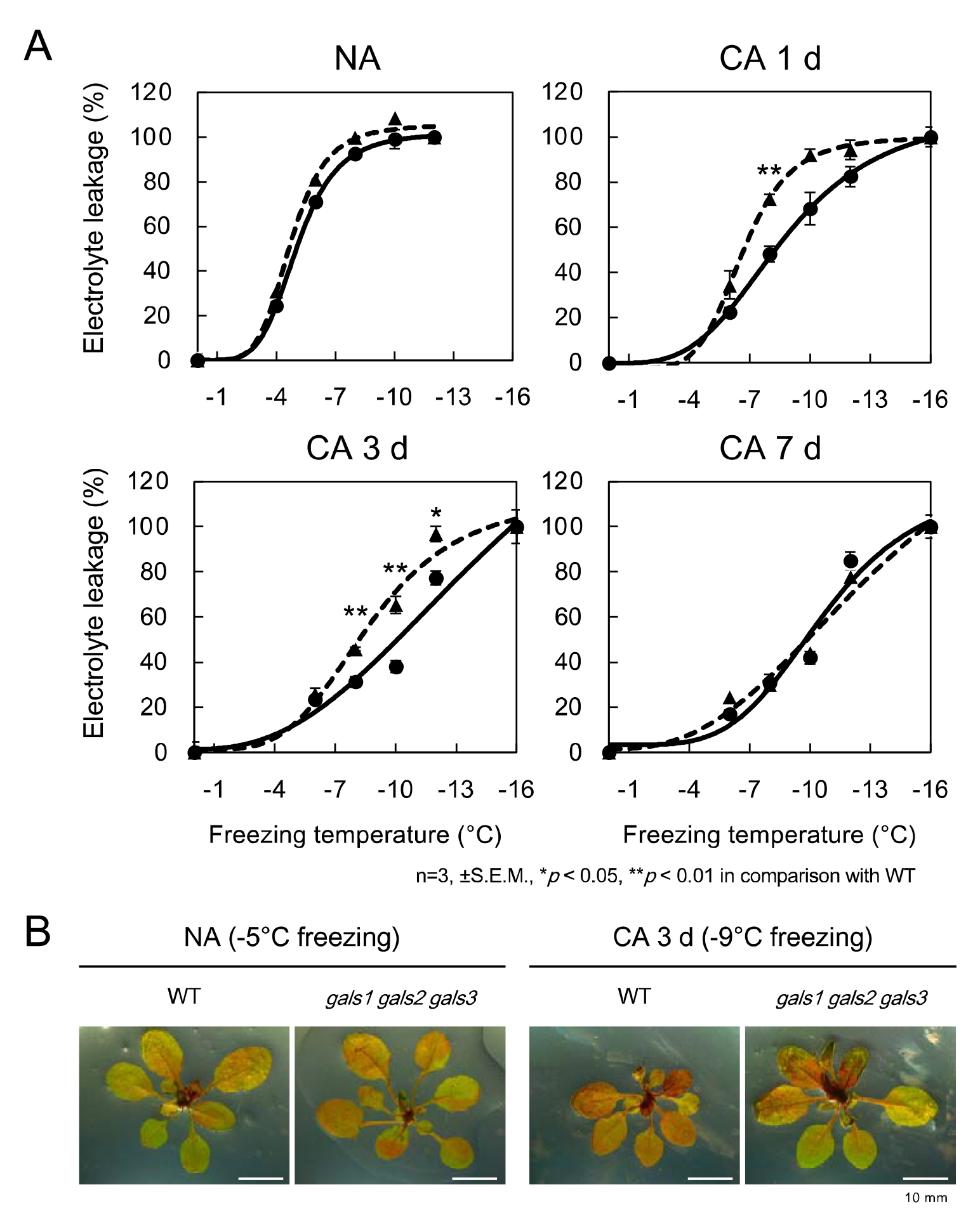
Decreased pectic galactan levels increase susceptibility to freezing after cold acclimation. (A) Changes in freezing tolerance of wild type (WT) and *gals1 gasl2 gals3* during the cold acclimation (CA) process. Electrolyte leakage assay was used to determine the freezing injury of plants grown on mineral wool at each freezing temperature. Solid lines indicate WT and dashed lines indicate *gals1 gasl2 gals3*. Error bars indicate ± S.E.M. (n = 3). Significant differences (Student’s *t*-test) between WT and *gals1 gasl2 gals3* are indicated by asterisks above the lines (\**p* < 0.05, \*\**p* < 0.01). (B) To examine tissue-specific viability of plants exposed to various freezing temperatures, surviving parts (red) were stained with 2,3,5-triphenyltetrazolium chloride (TTC) reagents. Scale bars, 10 mm.

**Figure 4.**
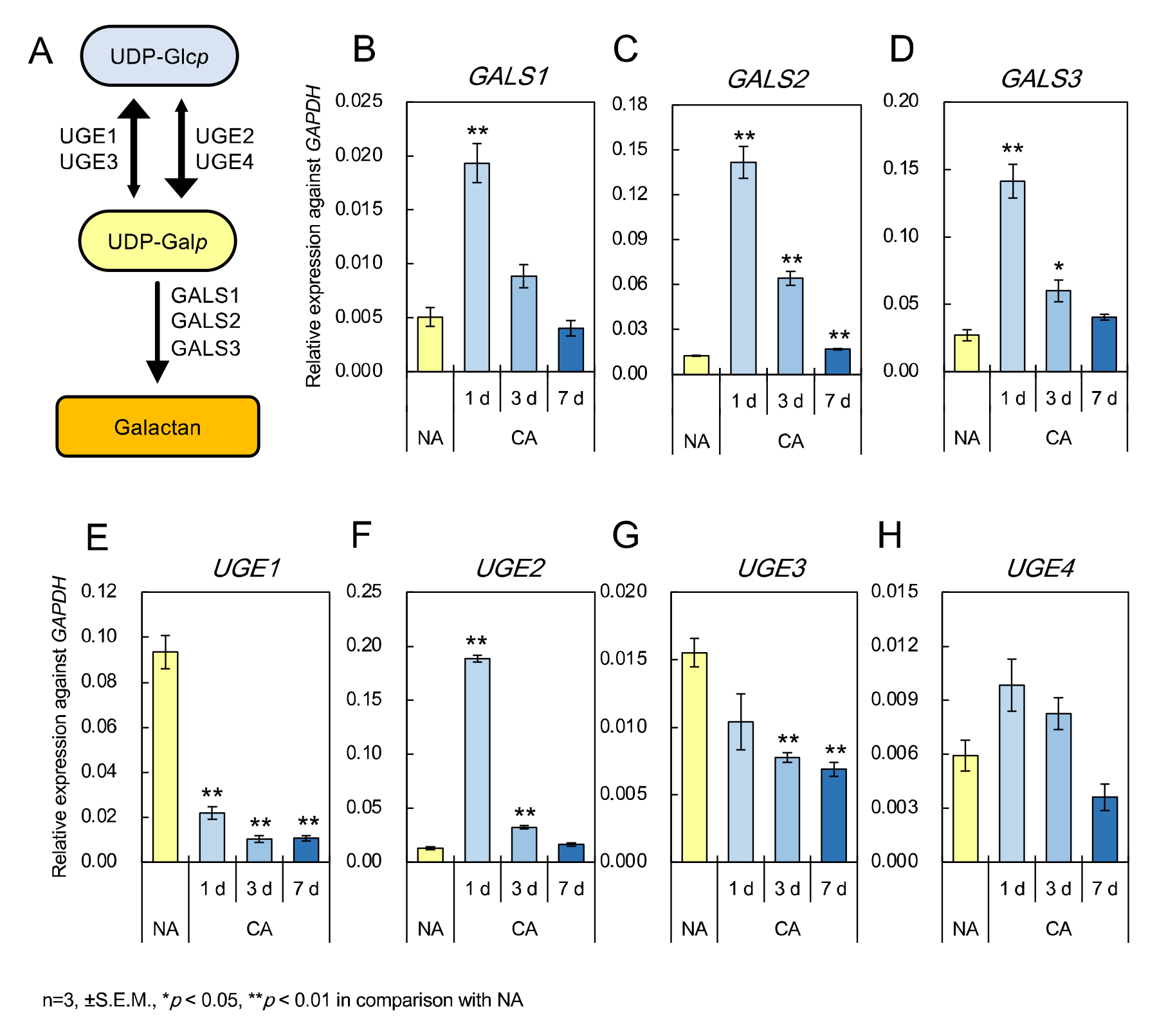
Changes in the expression of pectic galactan synthesis-related genes in *Arabidopsis thaliana* during cold acclimation. (A) Scheme for the synthesis of pectic galactan in Arabidopsis. UDP-Gal*p* is converted from UDP-Glc*p* by the contribution of UGE2 and UGE4. UGE1 and UGE3 predominantly perform the reverse reaction. Pectic galactan is synthesized by GALS using UDP-Gal*p* as a substrate. (B-D) Expression patterns of three *GALS* genes encoded by the Arabidopsis genome during the cold acclimation (CA) process. (E-H) Expression patterns of four *UGE* genes in Arabidopsis during CA. (B-H) Values represent the expression level relative to that of *GAPDH*. Error bars indicate ± S.E.M. (n = 3). Significant differences (Dunnett’s test) between non-acclimation (NA) and other treatment samples are indicated by asterisks above the bars (\**p* < 0.05, \*\**p* < 0.01).

Analysis of the *cis*-element in their promoter regions in Arabidopsis by plant promoter database ver. 3.0 (PPDB, https://ppdb.agr.gifu-u.ac.jp/ppdb, 32) predicted that *UGE2* and *UGE4* have an abscisic acid (ABA)-responsive *cis*-element in their promoter regions (AtREG468 and 379 for *UGE2* and AtREG608, 367 and 450 for *UGE4*, Figs. S10). Since ABA-responsive *cis*-elements are present in many cold-responsive genes, it is possible that the expression of the above genes may be altered by CA (8).

We therefore examined the expression of genes involved in galactan synthesis in Arabidopsis by RNA sequencing (RNAseq) (Dataset S1). In addition to NA and CA plants, de-acclimated (DA) plants, which were dehardened by transfer to room temperature after CA, were also analyzed. According to Venn diagrams (Fig. S11) and principal component analysis (PCA, Fig. S12) of RNAseq data, gene expression was altered by CA and partly returned to NA levels by DA treatment. Low temperature response markers including master transcription factors of the CA responses in plants (*CBFs/DREBs*) and their downstream genes (*CORs*, *GolSs*, and *RD29s*) were all increased in CA and decreased in DA (Fig. S13).

RNAseq data showed that the three *GALSs*, *UGE2,* and *UGE4* were all induced in CA in conjunction with the changes in CA response markers, while expression of *UGE1* and *UGE3* was repressed in CA (Fig. S14). Expression of the major *β-Galactosidase* (*BGAL*) genes potentially involved in pectic galactan degradation (e.g. *BGAL1* and *BGAL6*) (40–42) was decreased in CA (Fig. S15). On the other hand, expression of many of the galactan biosynthesis-related genes as well as cold response markers returned to NA levels in DA (Fig. S13 and S14) and appeared to be linked to changes in freezing tolerance during CA and DA. We thus performed RT-qPCR to investigate the temporal changes in gene expression of pectic galactan biosynthesis-related genes during the CA process. The results showed that the expression of the three *GALSs* (Fig. 4*B*-*H*) and those of *UGE2* and *UGE4* (Fig. 4*E*-*H*) were strongly induced at the initial stage of CA. By contrast, the expression levels of *UGE1* and *UGE3*, which are less involved in galactan synthesis, decreased during CA. These results imply an increase in the UGP-Gal pool and enhancement of pectic galactan synthesis by increased expression of *GALSs* and several *UGEs* and a reduction in pectic galactan turnover.

### Accumulation of pectic galactan affects tissue mechanical properties

What properties of the cell wall are affected by the accumulation of pectic galactan and influence plant freezing tolerance? There are two ways that cell wall remodeling may impact freezing tolerance: by increasing the hydration capacity of the cell wall (thereby reducing “free” water molecules in the wall) and/or by increasing stiffness of the cell wall (11, 13).

If cell wall remodeling for CA is intended to alter the interaction between the cell wall and water molecules, this presumably works by increasing the proportion of weakly cell wall-bound water (water molecules resistant to freezing) in the cell wall. This would decrease the proportion of free water, thereby alleviating the effects of freezing (13). To investigate this possibility, the heat flux of the leaves while temperature decreased was measured using differential scanning calorimetry (DSC, Fig. S16). The freezing point appeared to decrease, and the proportion of bound water appeared to increase during CA, in both WT and *gals1 gals2 gals3*, and there was no difference between WT and the triple mutant. This suggests that the moderation of the effect of extracellular freezing by CA is not due to a relative increase in cell wall bound water caused by an increase in cell wall pectic galactan during CA.

To investigate the involvement of pectic galactan in cell wall mechanical properties, we evaluated the physical properties of plant tissues by tensile strength testing (Fig. 5). In this analysis, the tissue was immersed in ethanol to counter the effects of turgor pressure and tissue pH on the physical properties of the tissue prior to measuring the extensibility and breaking force of the cell wall. WT plants showed a decrease in cell wall extensibility and an increase in rigidity as CA progressed. By contrast, both parameters of cell wall mechanical properties in *gals1 gals2 gals3* did not change significantly in CA. This indicates that pectic galactan accumulation is a key contributor to the mechanical changes of the cell wall associated with CA.

**Figure 5.**
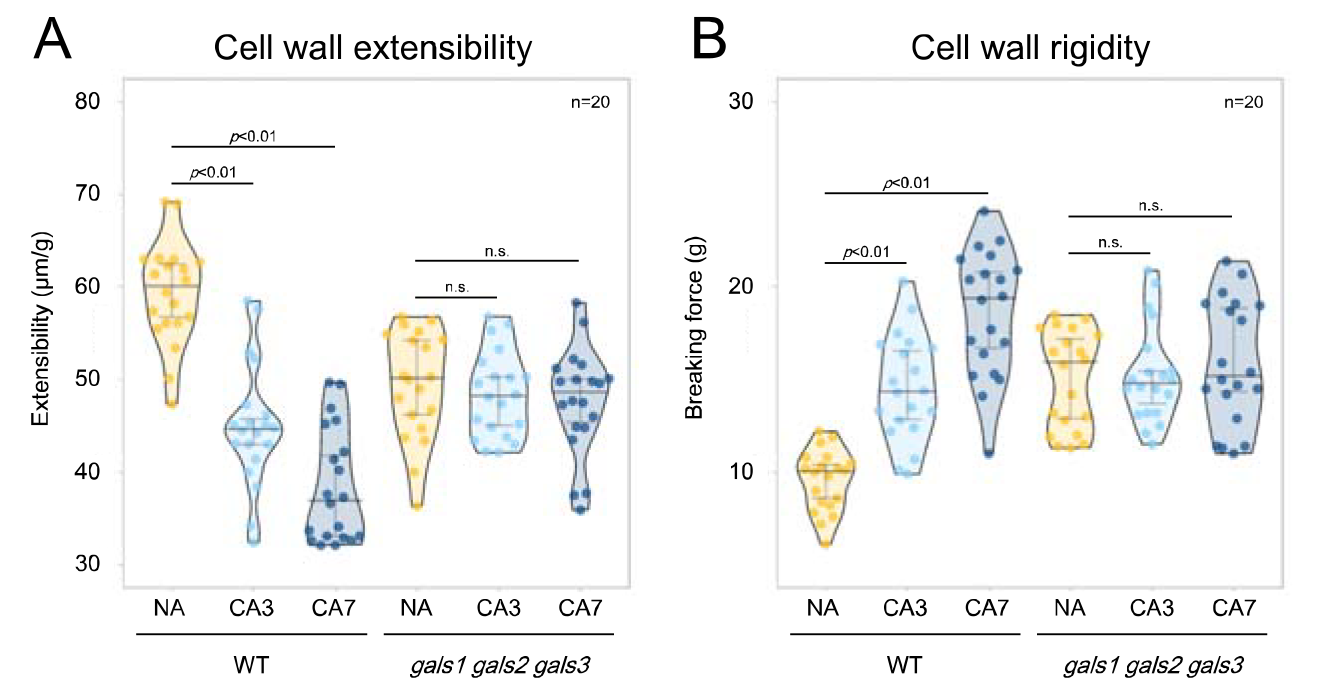
Pectic galactan levels affect stiffness of the cell wall during cold acclimation. Changes in the mechanical properties of leaf cell walls during cold acclimation (CA) were quantified by tensile testing. To measure the physical properties of plant cell walls, strips of tissue were immersed in ethanol to eliminate turgor pressure. From the values obtained, extensibility (A) and rigidity (B) were calculated and plotted on a graph (n = 20). Horizontal lines display the median value, and vertical bars show the 95% confidence interval. Significant differences at *p* < 0.05 and *p* < 0.01 (Dunnett’s test) between non-acclimation (NA) and other treatment samples are indicated above the bars (n.s., not significant).

## Discussion

Molecular mechanisms contributing to freezing adaptation common to temperate herbaceous plants are still largely unknown, except for the accumulation of soluble sugars and cold-responsive proteins and changes of membrane composition. There appears to be a common cell wall response pattern to CA, at least in several dicotyledonous plants. In this study, we provided evidence for the contribution of pectic galactan in CA. Accumulation of pectic galactan during the CA process was inferred in spinach, Japanese mustard spinach, and crown daisy in this study, in addition to *Arabidopsis thaliana*. In the peas used in this study, the relative amount of Gal in the pectin fraction did not appear to increase after 7 days of CA treatment, but its absolute amount increased. In a previous study, accumulation of pectic galactan by CA for 10 d was reported in a freezing-tolerant pea cultivar, which was not found in a freezing-susceptible cultivar (31). The findings of this study and previous work support the hypothesis that pectic galactan accumulation is an almost ubiquitous cold response of plants adapting to a freezing environment, like the accumulation of soluble sugars and cold-responsive proteins.

Enzymes of the GT92 family, to which GALSs belong, have only been identified in land plants and some animals, but not in green algae (27). The plant GT92 family is specialized for β-1,4-galactan synthesis. However, monocots, *Physcomitrium patens* and *Selaginella moellendorffii* have no or very little pectic β-1,4-galactan(43, 44). This suggests that the presence of pectic galactan in dicotyledons may have implications specific to dicots. The clades to which AtGALS2 and AtGALS3 belong are relatively distant from that of AtGALS1 and the clades of other plant enzymes and occur only in dicot plants including *Populus trichocarpa* (poplar). This suggests that these GALSs may play an important role in the process of acquiring freezing tolerance in many dicots. However, the presence or absence of these genes does not appear to be associated with freezing tolerance in general, since the tomato *Solanum lycopersicum*, which has low chilling-tolerance, also has orthologs of these GALSs. This may be because freezing tolerance is determined by the interaction of various factors (45).

The expression of *GALS* and *UGE* genes was rapidly altered by CA. Expression of *GALS1* has been shown to be induced by salt stress *via* BARLEY B RECOMBINANT/BASIC PENTACYSTEINE (BPC) 1/BPC2 transcription factor (29). It is unclear whether induction of *GALS1* by CA occurs by the same mechanism. A yeast one-hybrid assay also suggests that CBF2, a known master transcription factor of CA binds to the promoter region of *GALS1* (29). The relatively early induction of *GALS1* expression in response to CA would be consistent with regulation by CBF. Since ABA is the major plant hormone induced in CA, the induction of *UGE* may be regulated by ABA accumulation under CA conditions. In addition, *UGE2* has been shown to be down-regulated in a loss-of-function mutant of three *CBF* genes (46) and *GALS2* and *UGE2* are induced by ABA (47). Therefore, it will be necessary in the future to unravel the interrelationships between the regulation of β-1,4-galactan synthesis *via* gene expression in *GALSs* and *UGEs* and the CA response system from CBF and ABA.

This study supports the hypothesis that increased *GALSs* expression and pectic galactan contribute to changes in cell wall mechanical properties and freezing tolerance during CA in *Arabidopsis thaliana*. The question of whether tissue stiffness directly affects freezing tolerance remains. Previous studies have pointed out CA-induced increases in cell wall stiffness in various plant species *via* measurements of the elastic modulus (12, 17, 48–50). High cell wall stiffness protects the cell interior from external ice crystal expansion (11, 51). Related to this, in xylem parenchyma cells of trees, sufficiently stiff cell walls are believed to contribute to deep supercooling capacity (52, 53). Comparative studies of tissues in various plants have indicated a possible correlation between the elastic modulus and the degree of freeze-induced dehydration (54). Rigid cell walls are believed to exert a negative turgor pressure on living cells undergoing freeze-dehydration, thereby reducing the extent of freezing dehydration and eventually alleviating freezing injury (11, 54). By contrast, it has also been proposed that the microcapillary structure of the cell wall affects the propagation of ice in the tissue. In artificial materials that mimic a cell wall, microcapillaries smaller than 100 nm hinder ice propagation (51, 55). In some plants, e.g., *Allium fistulosum*, cell wall permeability is remarkably reduced with increased freezing tolerance during CA (21), and changes in cell wall porosity appear to be linked to cellular freezing tolerance(56). It is probable that galactan accumulation changes the mobility of solutes in the microcapillary structure as well as the mechanical strength of the cell wall (57).

In this study, freezing tolerance was restored to WT levels after CA for 7 d even in mutants that were deficient in pectic galactan. The triple *gals1 gals2 gals3* mutant does not accumulate pectic galactan in CA like WT, but instead has increased Ara and GalA. The higher amount of arabinan and HG in late CA in *gals1 gals2 gals3* compared to WT may involve a compensation among cell wall components. Interestingly, pea has a much higher percentage of Ara in the pectin fraction than other species (Fig. 1*B*), suggesting that it has a higher amount of pectic arabinan, which, together with galactan, composes the neutral side chains of RG-I. The high relative amount of pectic arabinan may be the key to the high freezing tolerance of pea, which accumulate less pectic galactan. It has been proposed that in the resurrection plant *Myrothamnus flabellifolius*, pectic arabinan accumulation contributes to the maintenance of cell wall flexibility and rapid recovery from desiccation (58–60). This may be related to the question considered here, since under freezing stress, plants are often subjected to both dehydration and hyperosmotic stress due to extracellular freezing. Furthermore, although pectic arabinan is considered important for the mechanical properties of plant tissues, including stomata (24–26), the accumulation of arabinan and HG in the *gals1 gals2 gals3* mutant observed in CA for 7 d does not appear to complement the changes in mechanical properties caused by the reduction of pectic galactan. This indicates that pectic galactan and arabinan are not exact functional complements, but rather achieve similar purposes in the coordinated work of a complex cell wall network.

## Materials and Methods

### Plant materials and growth conditions

Arabidopsis (*Arabidopsis thaliana*) Columbia-0 seedlings were grown on Murashige-Skoog (MS) agar medium at 22°C under light and dark conditions (16/8 h light/dark, 120 µmol m^-2^s^-1^) for 7 days. The plants were further grown for an additional 10 to 14 days on mineral wool (Nittobo, Japan) under the same temperature and light conditions and designated as NA (non-acclimated) plants. For cold treatment, NA plants were grown at 4°C under light and dark conditions (12/12 h light/ dark, 120 µmol m^-2^ s^-1^) for up to 7 days (CA). The *gals1 gals2 gal3* mutant (SALK_016687, SALK_121802, WiscDsLox377-380G11, respectively) was provided by Henrik V. Scheller (28). Single and double mutants were generated by crossing with WT. Seeds of pea (*Pisum sativum* ‘753’), spinach (*Spinacia oleracea* ‘Progress’), Japanese mustard spinach (*Brassica rapa* var. *Perviridis* ‘Hamatsuduki’) and crown daisy (*Glebionis coronaria* ‘Satoakira’) purchased from Sakata seed corporation (Kanagawa, Japan) were sown on culture soil, and grown and cold-acclimated under the same conditions as *Arabidopsis thaliana*.

### Evaluation of freezing tolerance

Electrolyte leakage measurements were based on the methods of Thalhammer et al. (61) and Uemura et al. (62). The above-ground parts of plants were placed in test tubes with 60 µL of water. These were incubated at −2°C for 30 min with a programmable freezer (NCB-3300, EYELA, Japan), followed by the addition of ice grains, and incubation for 60 min. The temperature was then lowered at a rate of 2°C per h. Samples were taken out of the bath at designated temperatures and thawed at 4°C for 12 h in the dark. The samples were shaken for 2 h in 3 mL water. Electrical conductivity was measured with an EC meter (LAQUAtwin EC-33, Horiba, Japan), the sample was boiled for 15 min, and conductivity was measured again. The degree of freezing injury was determined by comparison of the conductivity before and after boiling.

Visualization of freezing injury with TTC assay was conducted as described in Livingston et al. (63) with a slight modification. Plants frozen in the same manner as above were taken out at the specified temperature and soaked in a solution composed of 0.5% (w/v) TTC and 50 mM HEPES (pH 7.3). Plants were incubated at room temperature for 12 h in the dark. Pictures were taken with a Stereomicroscope (S9i, Leica Microsystems, Germany).

### Cell wall fractionation and determination of monosaccharide composition

Fractionation and compositional analysis of the cell wall was performed as in to our previous study (14). Aboveground parts of the plants (100 to 200 mg) were ground in 1 mL water. The homogenized sample was centrifuged at 21,500 *g* for 5 min at 4°C. The following centrifugal steps were performed in the same manner each time. The supernatant was collected as the soluble fraction. The remaining precipitate was suspended in 80% ethanol, boiled for 2 min, and centrifuged. The pellet was suspended in amylase reaction solution (25 mM Mops-KOH, pH 6.5, 0.04 units (20 units/mL) α-amylase) and incubated at 37°C for 2 h. After centrifugation, the supernatant was collected. The precipitate was suspended in water, boiled for 10 min, and centrifuged. The supernatant was collected as the hot water fraction. This step was repeated twice. The same procedure was followed while replacing the solutions in the following order: 50 mM EDTA (pH 8.0)/50 mM sodium phosphate buffer and 4 M KOH supplemented with 10.6 mM NaBH_4_, as the EDTA and KOH fractions, respectively. The KOH fraction was neutralized by the addition of glacial acetic acid. The precipitate was washed with water, ethanol, and diethyl ether and air-dried overnight. The resultant powder was hydrolyzed in 72% sulfuric acid for 1 h, then in 8% sulfuric acid for 4 h and collected as the cellulose fraction. The total sugar content was determined by the phenol-sulfuric acid method (64).

Pectin-rich EDTA fractions were dialyzed against water for 2 days and lyophilized. Samples containing 50 µg of sugar were hydrolyzed in 2 N trifluoroacetic acid (TFA) for 60 min at 120°C. The hydrolyzed samples were dried by Speed Vac (TAITEC VC-36R, Japan) and redissolved in water. After repeating this step three times, samples were dissolved in 250 µL of water. 50 µL of sample was subjected to high-performance anion exchange chromatography-pulsed amperometric detection (HPAEC-PAD) with the ICS −5000⁺series system equipped with a CarboPac PA-1 column (Thermo Fisher Scientific, USA). Elution was carried out using water, 0.1 M sodium hydroxide with 0.5 M sodium acetate at a flow rate of 1.0 mL min^-1^ as described previously (65).

### Determination of pectic galactan content using β-1,4-galactanase

A 10 µL suspension of dialyzed pectin-rich EDTA fraction (50 µg sugar content) was added to 90 µL enzyme solution containing 50 mM sodium-phosphate buffer (pH4.5) and endo-β-1,4-galactanase from *Aspergillus niger* (1.3 units, Megazyme, Ireland) and incubated at 37°C for 12 h. After boiling for 5 min and centrifuging, maltopentaose was added to the supernatant as an internal standard, and free sugars were labeled with ABEE according to a previous study (66) and analyzed by HPLC.

### Immunohistochemical staining of pectic galactan in leaf tissues

The fifth youngest leaf of the plant was cut and soaked in fixing solution containing 4% (v/v) paraformaldehyde and 0.025% (v/v) glutaraldehyde. Samples were vacuum-infiltrated and incubated for 12 h. After washing the samples with phosphate-buffered saline (PBS), they were immersed in 30% (v/v) ethanol and allowed to stand for 6 h. This was repeated, successively replacing the solution with a 50%, 70%, 80%, 90%, 100%, and 100% ethanol. Similar steps were performed by successively replacing ethanol with a mixture of ethanol and Technovit 7100 resin (Kulzer Technik, Germany) containing 1% (w/v) hardener I in the following proportions: 5:1, 3:1, 2:3, 1:5, 0:1 and 0:1. Finally, this was replaced with Technovit 7100 containing 10% (v/v) Hardener II and allowed to cure for 12 h at 60°C. Ten µm thick sections were cut from the resin in which the samples were embedded and placed on glass slides. Sections were treated with blocking solution containing 3% (w/v) skim milk in PBS for 30 min and then with LM5 antibody (Kerafast, USA) diluted 5-fold in blocking solution for 1 h. The same treatment was performed with the antibody replaced by water as a negative control. The sections were then washed three times with PBS, treated with secondary antibody (Alexa Fluor 488, Thermo Fisher Scientific, USA) for 1 h, and washed twice with PBS. After staining with Calcofluor (Sigma-aldrich, USA), the sections were observed under a confocal microscope at excitation wavelengths of 405 nm and 473 nm (FV1000, Olympus, Japan).

### RNA extraction, RT-qPCR analysis, and RNAseq

Total RNA was extracted using Isogen (Nippon Gene, Japan) according to manufacturer′s instructions. RNA samples (1 µg) were converted to cDNA with oligo dT primers according to the instructions of the PrimeScript RT reagent Kit with gDNA Eraser (Takara, Japan). Quantitative PCR was performed in accordance with manufacturer’s instructions for THUNDERBIRD SYBR qPCR Mix (TOYOBO, Japan) using the StepOne realtime PCR System (Thermo Fisher Scientific, USA). *GAPDH* (*At1g13440*) was used as the internal standard gene. All primers used in the experiment are listed in Table S1. For RNAseq, the obtained RNA (10 µg) was treated with DNaseI (Invitrogen, USA) and purified with RNeasyPlant mini kit (Qiagen, Germany) in accordance with manufacturer′s instructions. RNA-Seq library construction, next generation sequencing, and bioinformatics analyses were run at GENEWIZ from Azenta Life Sciences (USA) using the poly-A selection method with DNBSEQ 2×150 bp (MGI Tech, Shenzhen, China).

### Measurement of mechanical properties of the cell wall

For measurement of mechanical properties of the cell wall, the fifth leaf of the plant was harvested and cut into strips of 3 mm width. The strips were immediately immersed in 80% ethanol and allowed to sit for at least 1 day. These samples were subjected to a tensile tester (Tensilon STB-1225S, A&D Co. Ltd., Tokyo, Japan) to determine extensibility and stiffness of the tissue cell wall. Each leaf strip was clamped at 1 mm intervals and stretched at a rate of 20 mm/min until it broke. The cell wall extensibility (strain/load in units of μm/g) was determined by measuring the load rate increase from 4 to 5 g.

### Statistical Analysis

Statistical significance of differences was determined by student’s *t*-test for two-group comparisons and Dunnett’s test or the Tukey-Kramer test for multiple comparisons at the *p* < 0.05 level using Microsoft Excel or the R software (ver. 4.2.1). The data from RNAseq analysis was Pareto-scaled for principal component analysis (PCA) using the *prcomp* function in R.

### Accession Numbers

The gene sequences and other information presented in this study are available in The Arabidopsis Information Resource (TAIR, https://www.arabidopsis.org) under the following accession numbers: GALS1 (AT2G33570), GALS2 (AT5G44670), GALS3 (AT4G20170), UGE1 (AT1G12780), UGE2 (AT4G23920), UGE3 (AT1G63180), UGE4 (AT1G64440), GAPDH (AT1G13440), CBF1 (AT4G25490), CBF2 (AT4G25470), CBF3 (AT4G25480), COR15A (AT2G42540), COR15B (AT2G42530), RD29A (AT5G52310), RD29B (AT5G52300), GolS1(AT2G47180), BGAL1 (AT3G13750), BGAL2 (AT3G52840), BGAL3 (AT4G36360), BGAL4 (AT5G56870), BGAL5 (AT1G45130), BGAL6 (AT5G63800), BGAL7 (AT5G20710), BGAL8 (AT2G28470), BGAL9 (AT2G32810), BGAL10 (AT5G63810), BGAL12 (AT4G26140), BGAL14 (AT4G38590), BGAL16 (AT1G77410), BGAL17 (AT1G7299).

## Acknowledgments

We are grateful to Prof. Matsuo Uemura (Iwate University) for his critical assistance in the preparation of the manuscript. This work was supported by funding from MEXT KAKENHI Grants-in-Aid for Scientific Research to D. T. [no. 20 K15494] and T. Kotake [nos. 16 K07391 and 19 K06702], by Grant-in-Aid for Scientific Research on Innovative Areas “Plant-Structure Optimization Strategy” to T. Kotake [no. 18H05495], by Ichimura Foundation for New Technology to D. T. [nos. 29-14 and 30-12] and by Kato Memorial Bioscience Foundation to D. T.

