## Supporting information for "Cell wall pectic β-1,4-galactan contributes to increased plant freezing tolerance induced by cold acclimation"

*Daisuke Takahashi

+81-48-858-3760

Graduate School of Science & Engineering, Saitama University, 255 Shimo-Okubo, Sakura-ku, Saitama 338-8570, Japan

**Materials and Methods**

**Glycosidic linkage analysis.** Glycosidic linkage analysis using gas liquid chromatography (GLC) was performed according to previous studies (1–3). Briefly, a 100 µg pectin fraction was aliquated and polysaccharides were pre-methylated according to the method by Pettolino et al (2). After adding 0.4 µg of *myo*-inositol to each sample as an internal standard, samples were hydrolyzed by 2 N TFA for 1 h at 121°C, derivatized to alditol acetates, and analyzed with a Shimadzu gas chromatograph GC-2014 fitted with a column (0.22 mm × 25 m) of BPX70 (Shimadzu, Kyoto, Japan). The 2,3,6-Me_3_-Gal (→4Gal*p*1→) and 2-Me-Ara (→3,5Ara*f*1→) peaks were identified using Galactan from lupin and Arabinan from sugar beet (Biocon, Nagoya, Japan) as standards.

**Phylogenetic tree analysis.** Amino acid sequences of GALS and UGE proteins for each plant species were obtained from the Phytozome Database (https://phytozome-next.jgi.doe.gov) (4).The amino acid sequences were aligned using ClustalW and then a phylogenetic tree was created using MEGA11 (5). The distance between each branch was determined using the neighbor-joining method with 1000 bootstrap samples. The amino acid sequences used in this study are listed in Table S2.

**Measurement of freezing behavior of the cell wall.** Leaves of the plants were collected and the cut surfaces of the main veins were dipped in water with a trace of Snomax powder (Snomax, USA), an ice nucleating protein to induce stable ice formation within the differential scanning calorimetry (DSC, DSC214 polyma, NETZSCH, Germany). After wiping excess water from the leaves, samples were sealed with aluminum rid. Empty pans were used as references. Both were set in the DSC, and the temperature was lowered to -80°C at a rate of 5°C/min. After maintaining at -80°C for 2 min, the temperature was raised to 25°C at a rate of 5°C/min. The dry weight of the AIR was calculated by punching a hole in the aluminum pan after the measurement and heating it at 65°C for 12 h, then re-measuring the weight. Peak areas of exotherm and endotherm were calculated from the obtained DSC curves with Proteus software (ver.7.0, NETZSCH). Calculation methods for unfrozen and intermediate water are in accordance with previous paper (6).

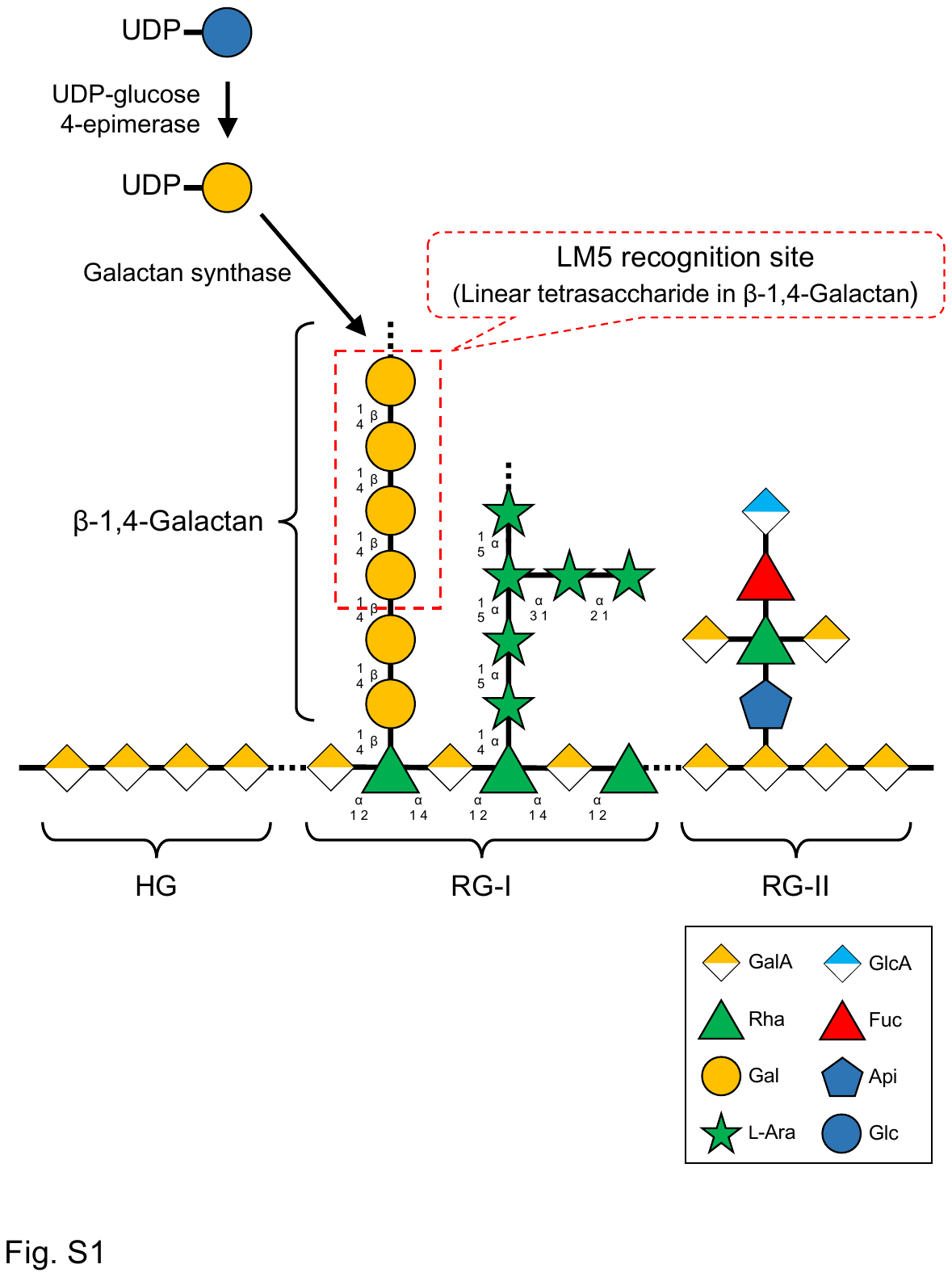
Fig. S1. Schematic diagram of β-1,4-galactan. The pectic galactan focused on in this study is a linear side chain of the pectin rhamnogalacturonan (RG)-I region composed of 1,4-linkages of galactose (Gal). GALACTAN SYNTHASE transfers Gal from the substrate, UDP-Gal, to pectic galactan. UDP-Gal is converted from UDP-Glc by UDP-glucose 4-epimerase. The LM5 antibody used in this study recognizes linear linear tetrasaccharide in β-1,4-galactan.

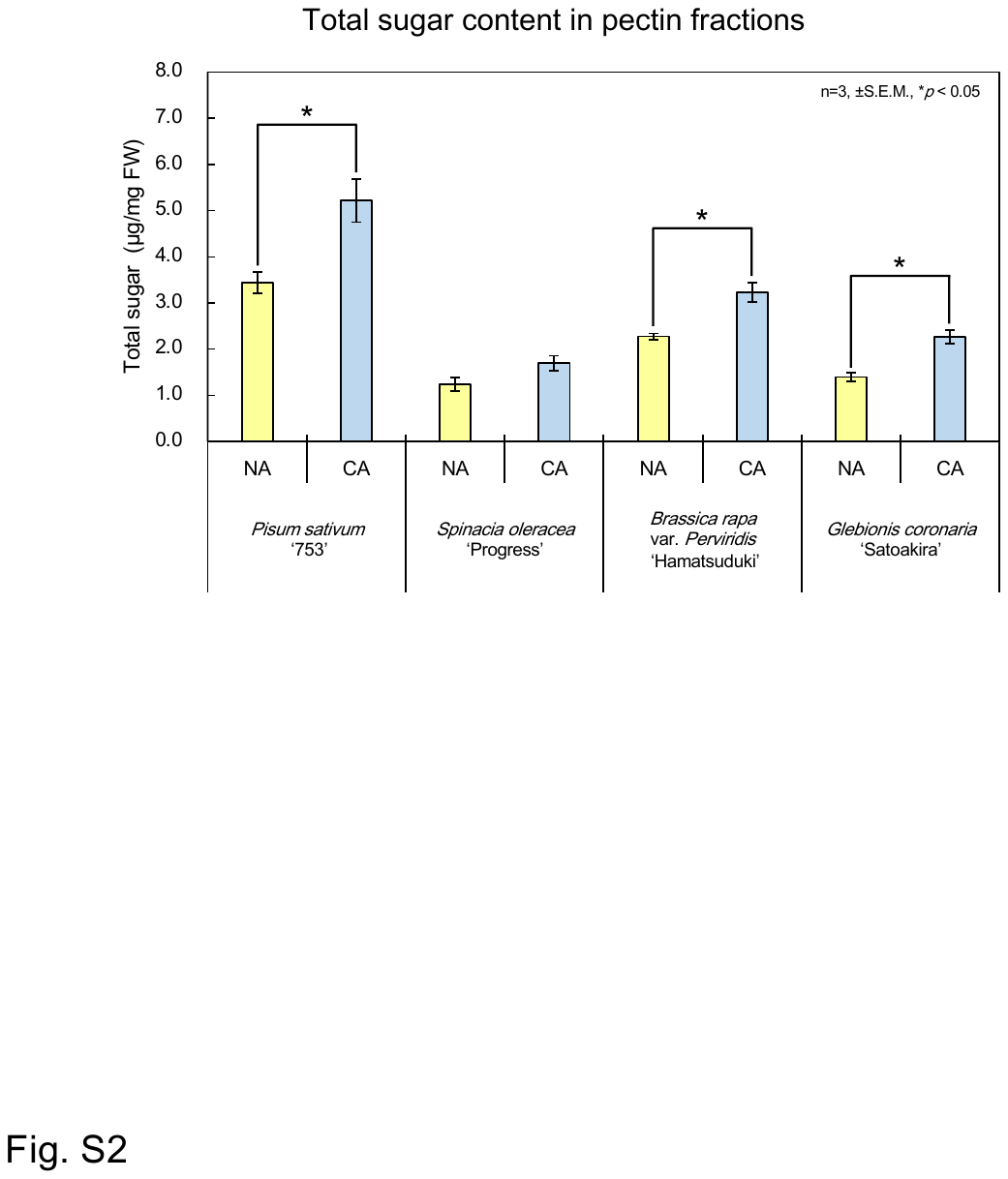
Fig. S2. Total sugar content of the pectin-rich raction in freezing tolerant vegetables. The pectin-rich fractions were extracted from cold non-acclimated (NA) and acclimated (CA) plants. The total sugar content in the fractions was determined by the phenol sulfuric method. Values represent the amount of sugar per fresh weight (FW). Statistically significant differences between NA and CA were determined with Student’s *t*-test (**p* < 0.05). Error bars indicate ± S.E.M. (n = 3).

Fig. S3. Monosaccharide composition in the pectin-rich fraction of four vegetables in non-acclimated (NA) and cold acclimated (CA) vegetables determined by HPAEC-PAD following acid hydrolysis. Values represent the amount of each sugar per fresh weight (FW). Statistically significant differences between NA and CA were determined with Student’s *t*-test (**p* < 0.05). Error bars indicate ± S.E.M. (n = 3). Fuc, fucose; Rha, rhamnose; Ara, arabinose; Gal, galactose; Glc, glucose; Man, mannose; Xyl, xylose; GalA, galacturonic acid; GlcA, glucuronic acid.

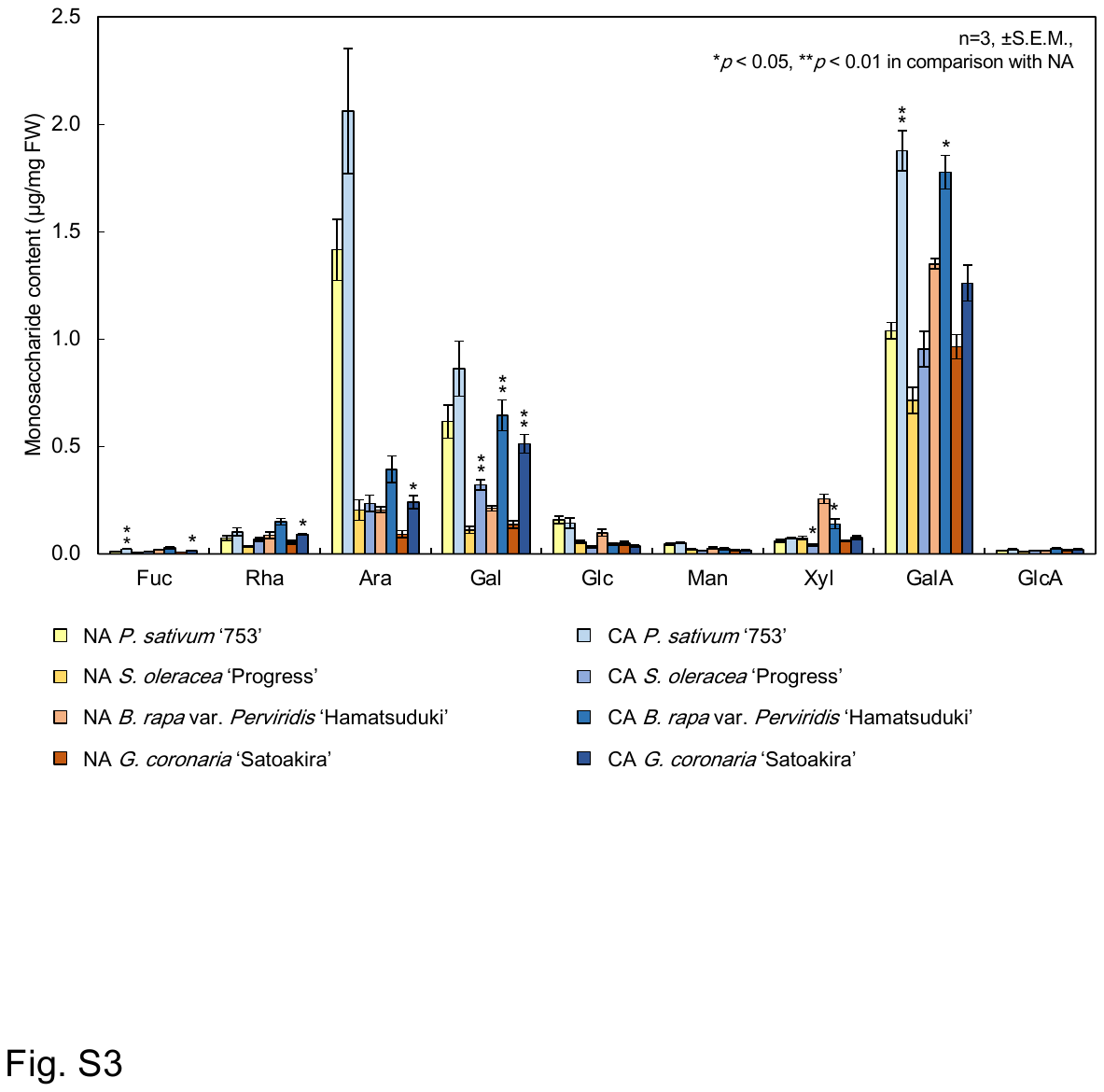

Fig. S4. Monosaccharide composition in the pectin-rich fraction extracted from non-acclimated (NA) and cold acclimated (CA) wild type (WT) and *gals1 gals2 gals3* determined by HPAEC-PAD following acid hydrolysis. The bar graph shows the percentage of each monosaccharide in pectin fractions. Statistically significant differences between WT and *gals1 gals2 gals3* were determined with Student’s *t*-test (**p* < 0.05). Error bars indicate ± S.E.M. (n = 3). Fuc, fucose; Rha, rhamnose; Ara, arabinose; Gal, galactose; Glc, glucose; Man, mannose; Xyl, xylose; GalA, galacturonic acid; GlcA, glucuronic acid.

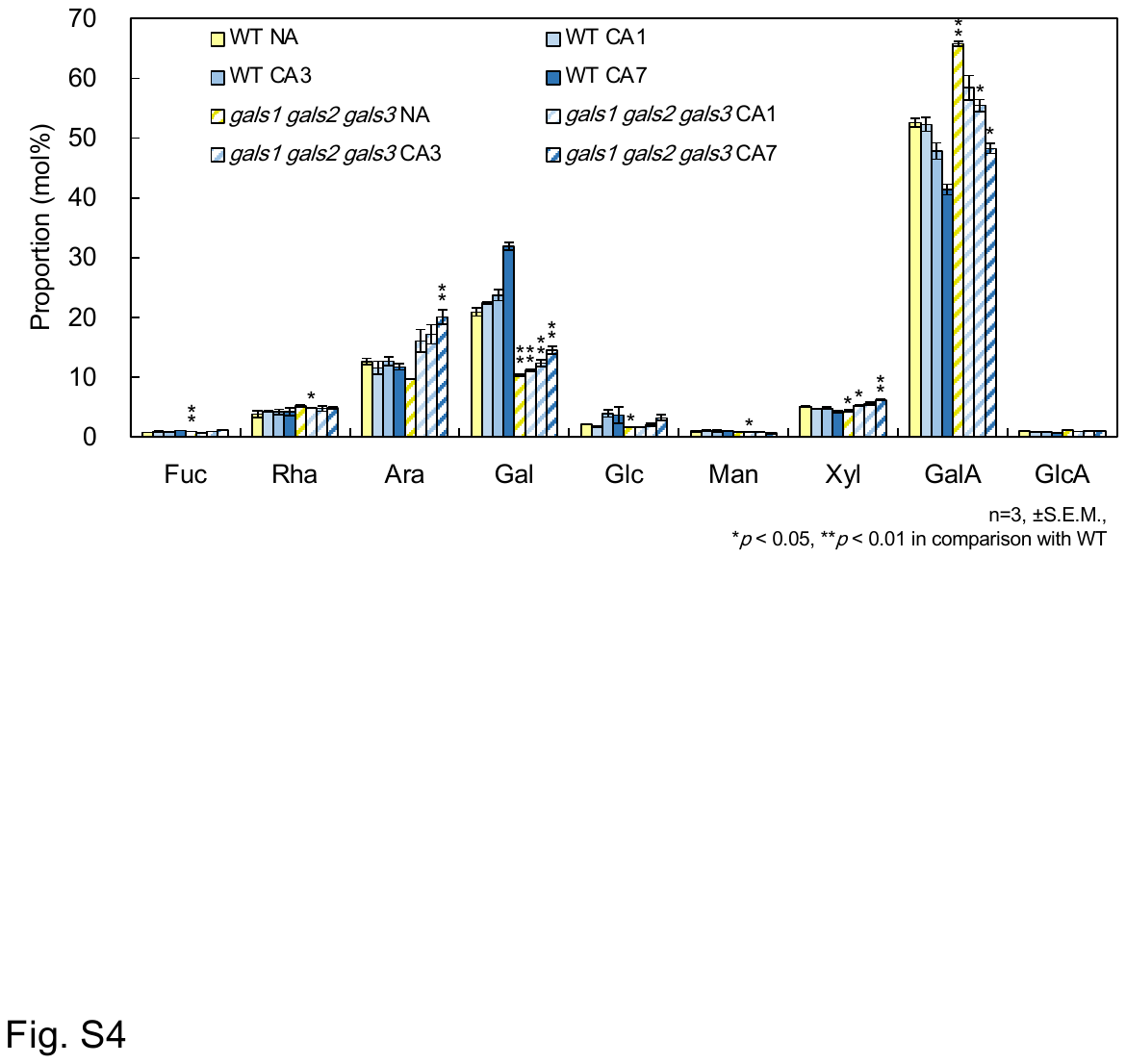

**Fig. S5.** Glycosidic linkage analysis by gas liquid chromatography (GLC). The methylated sugar peak derived from β-1,4-galactan in pectin fraction before and after cold acclimation for 3 d (CA3) of wild type (WT) and *gals1 gals2 gals3* mutant was identified using the β-1,4-galactan extracted from lupin as a standard.

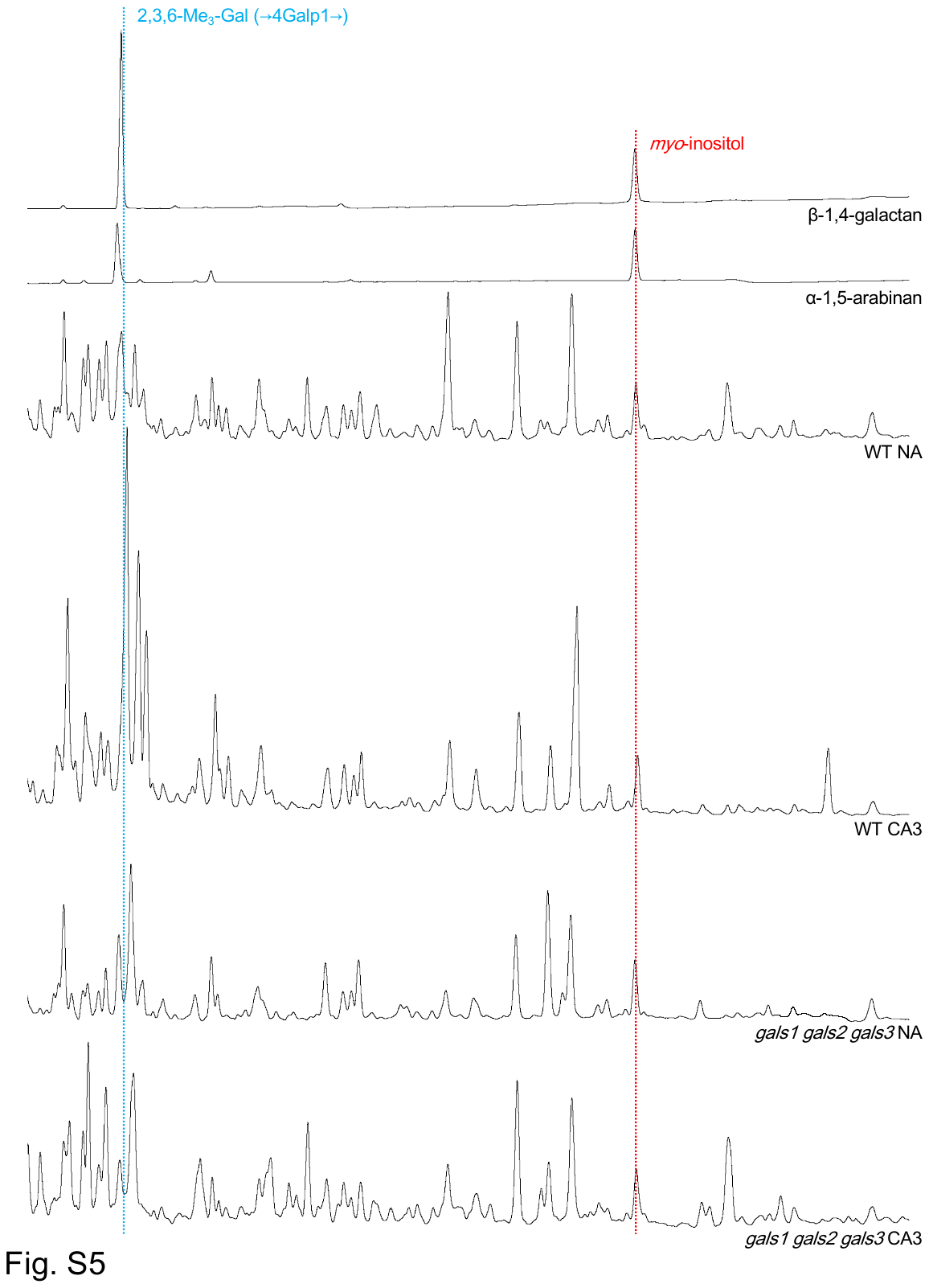

**Fig. S6.** Galactan amount in a series of *gals* single and multiple mutants. Pectin fractions extracted from non-acclimated (NA) and 3 d-cold acclimated (CA3) Arabidopsis were subjected to β-1,4-galactanase treatment, which specifically degrades pectic galactan, and the released galactobiose (Gal_2_) was detected by HPLC. The amount of galactan per fresh weight (FW) was estimated from the Gal_2_ peak area. Values represent the amount of Gal_2_ per FW. Statistically significant differences between wild type (WT) and others were determined with Student’s *t*-test (**p* < 0.05, ***p* < 0.01). Error bars indicate ± S.E.M. (n = 3).

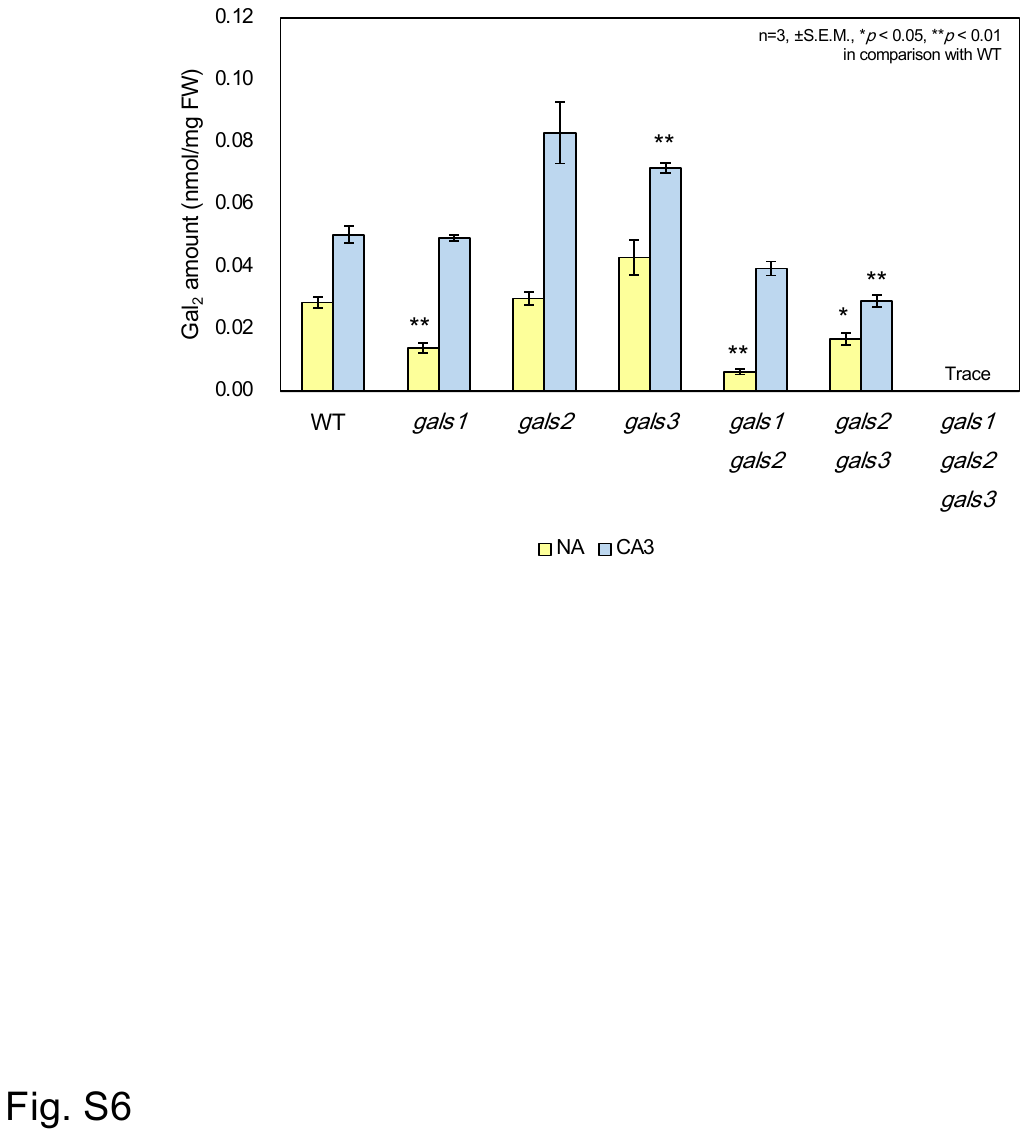

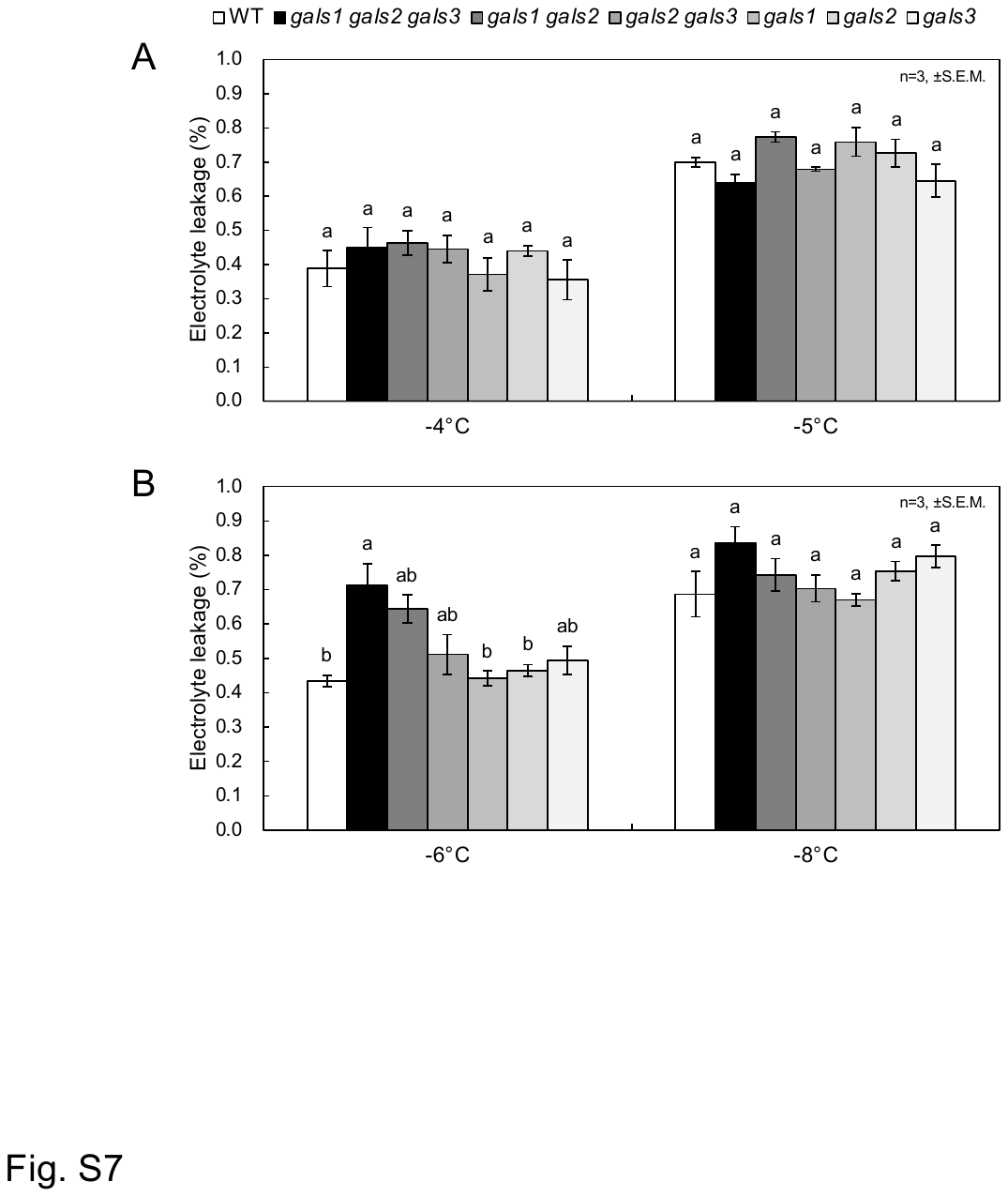
**Fig. S7.** Freezing tolerance of a series of *gals* single and multiple mutants in non-acclimation (NA) and cold acclimation (CA) for 3 d. NA plants grown on agar medium for 14 d (A) and NA plants followed by 3 days of CA treatment (CA3, B) were frozen at specific freezing temperatures. The electrical conductivity was measured before and after boiling, then the percentage of electrolyte leakage due to freezing injury was calculated. Statistically significant differences between genotypes were determined with the Tukey-Kramer test (*p* < 0.05) and are marked with lower case letters above the bars. Error bars indicate ± S.E.M. (n = 3).

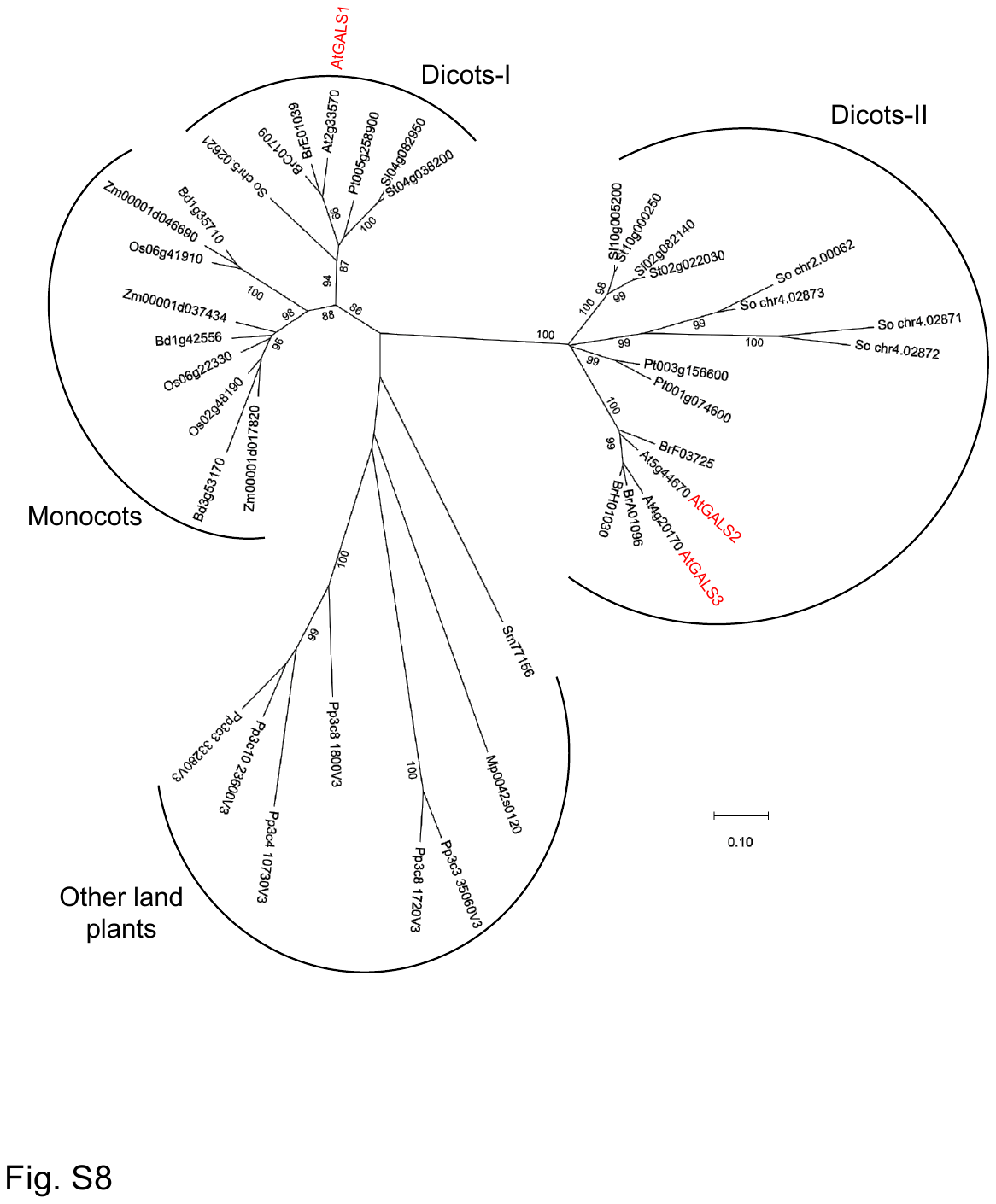
**Fig. S8.** Phylogenetic tree analysis of the *GALS* proteins. Amino acid sequences of various GALS homologue proteins were obtained from Phytozome. Only bootstrap values greater than 80% are shown. At, *Arabidopsis thaliana*; Bd, *Brachypodium distachyon*; Br, *Brassica rapa*; Mp, *Marchantia polymorpha*; Os, *Oryza sativa*; Pp, *Physcomitrium patens*; Pt, *Poplus trichocarpa*; Sm, *Selaginella moellendorffii*; Sl, *Solanum lycopersicum*; St, *Solanum tuberosum*; So, *Spinacia oleracea*; Zm, *Zea mays*.

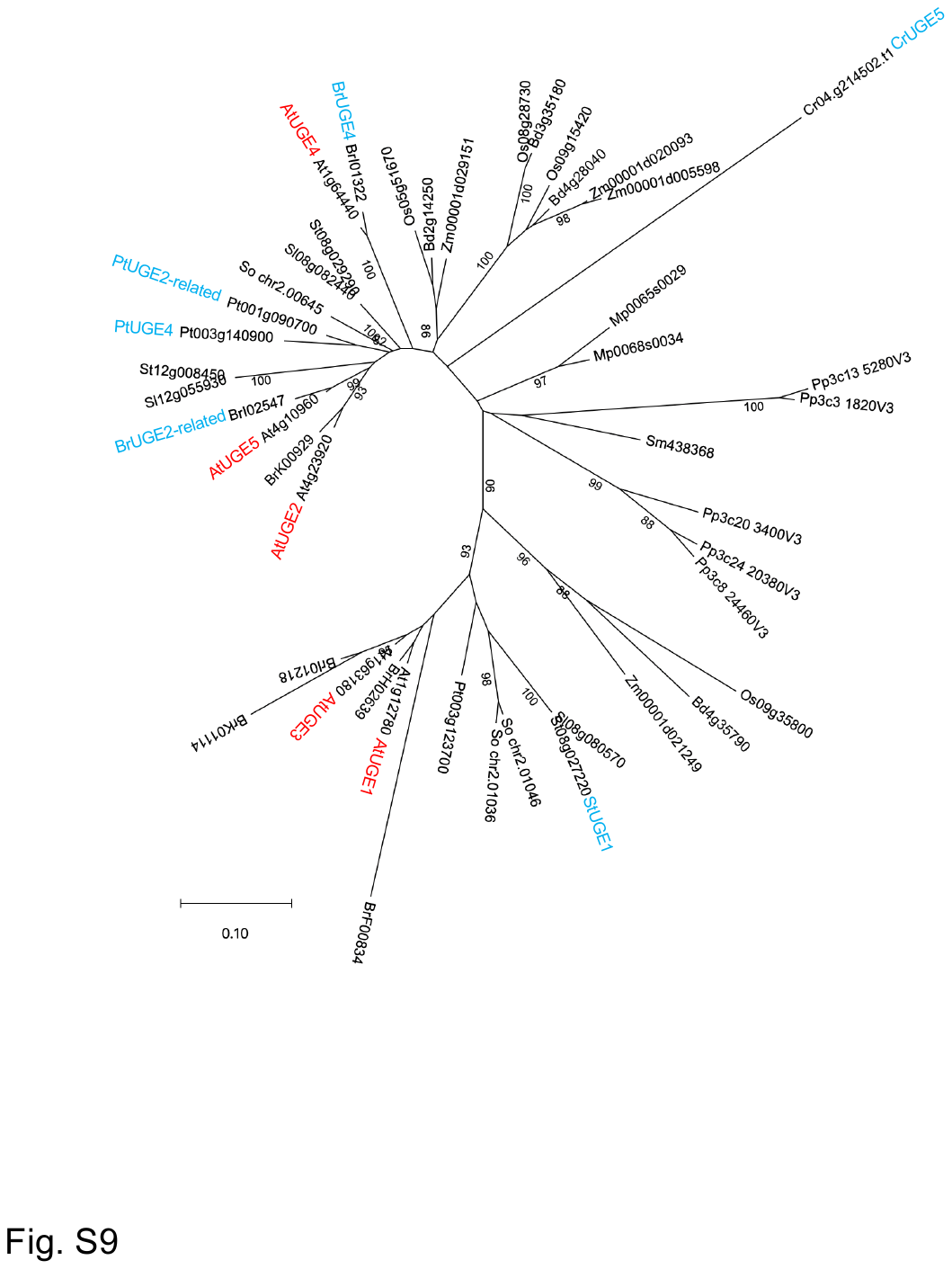
**Fig. S9.** Phylogenetic tree analysis of the *UGE* proteins. Amino acid sequences of various UGE homologue proteins were obtained from Phytozome. Only bootstrap values greater than 80% are shown. At, *Arabidopsis thaliana*; Bd, *Brachypodium distachyon*; Br, *Brassica rapa*; Cr, *Chlamydomonas reinhardii*; Mp, *Marchantia polymorpha*; Os, *Oryza sativa*; Pp, *Physcomitrium patens*; Pt, *Poplus trichocarpa*; Sm, *Selaginella moellendorffii*; Sl, *Solanum lycopersicum*; St, *Solanum tuberosum*; So, *Spinacia oleracea*; Zm, *Zea mays*.

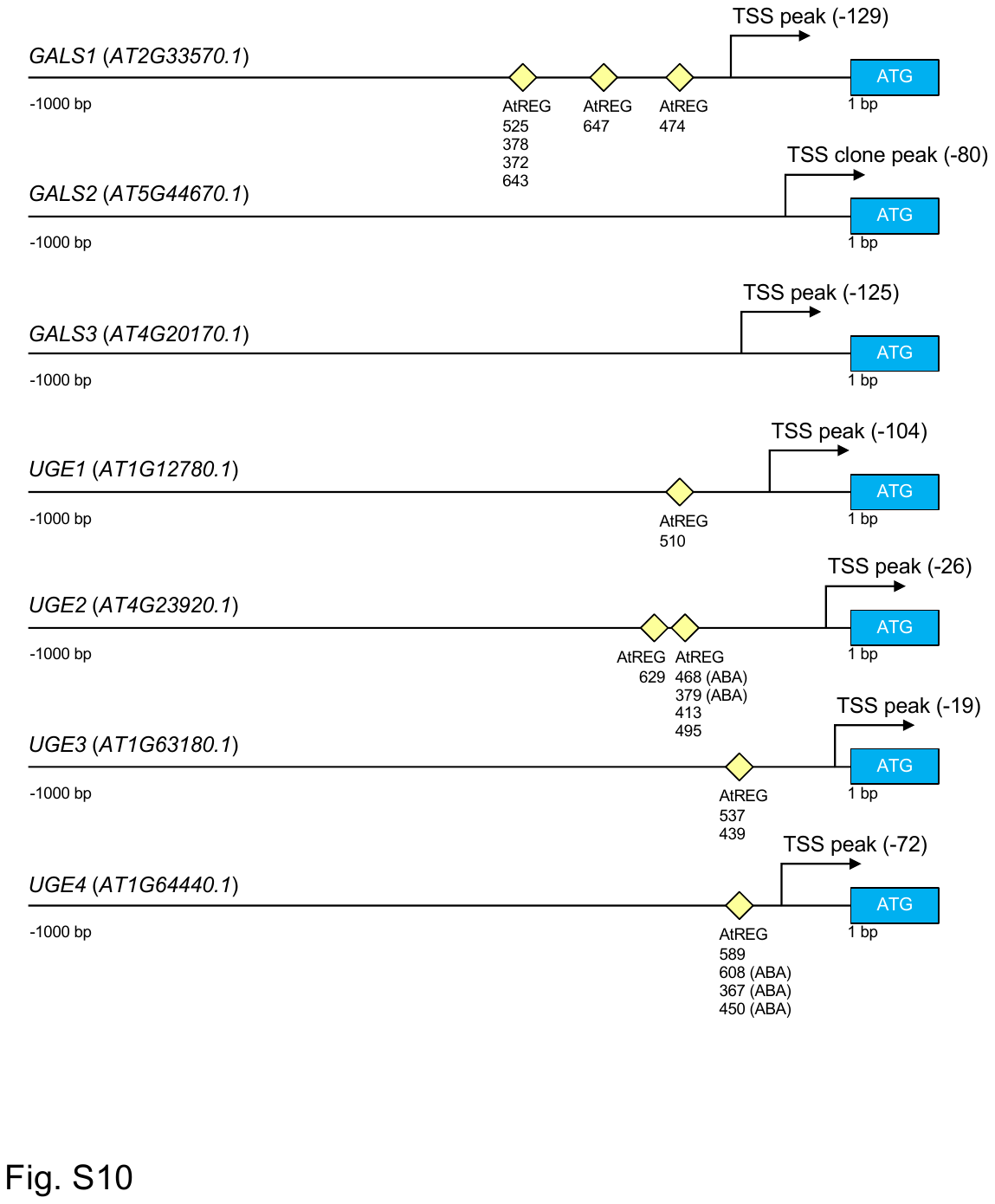
**Fig. S10.** *cis* elements in the promoter region of genes involved in galactan synthesis. *cis elements* of each gene were predicted by ppdb ver. 3.0 (https://ppdb.agr.gifu-u.ac.jp/ppdb/). The first codon is indicated by a blue box (ATG) and the transcription start site (TSS) is indicated by an arrow. Numbers represent base pairs from the TSS. The regulatory element groups (REGs) registered in the ppdb and their locations in the promoter region are indicated by yellow diamonds with accession numbers. Parentheses after the REG accession number indicate that it is a potential REG involved in the ABA response.

Fig. S11. Venn diagram classification of gene expression patterns in non-acclimation (NA), cold acclimation (CA), de-acclimation (DA) by RNAseq. (A) From RNAseq data (*SI Appendix*, Dataset S1), each gene was classified as having CA/NA and DA/CA fold change of 0.5 or less (*p* < 0.05 in Dunnett’s test), 2.0 (*p* < 0.05 in Dunnett’s test) or greater, or neither, and the number of genes in common in each comparison is represented in a Venn diagram. Expression of 1957 and 738 genes showed changes that are linked to freezing tolerance, such as an increase in CA and a decrease in DA, or a decrease in CA and an increase in DA, respectively. (B) From RNAseq data (*SI Appendix*, Dataset S1), genes involved in the cell wall in that gene ontology (GO) were extracted and classified as above. One sees that 115 genes were significantly altered upon CA and changed in a different direction from CA upon DA treatment.

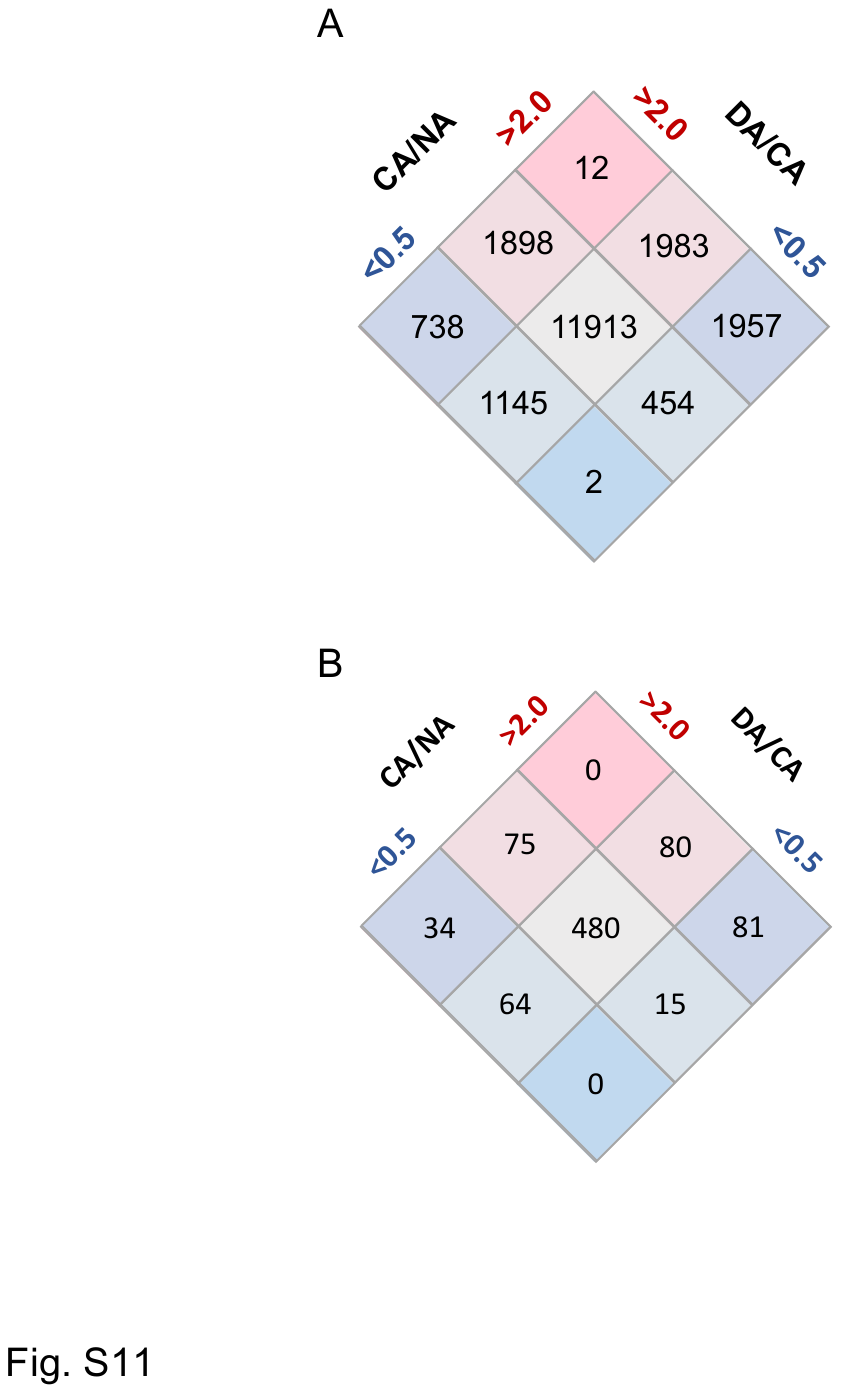

Fig. S12. Principal component analysis (PCA) of expression levels of genes in non-acclimation (NA), cold acclimation for 1 d (CA1) and de-acclimation for 1 d (DA1) retrieved from RNAseq data (*SI Appendix*, Dataset S1). Each point represents a sample. Score plots are shown for PC1 and PC2. The fraction of the total variance explained by each PC is indicated in parenthesis.

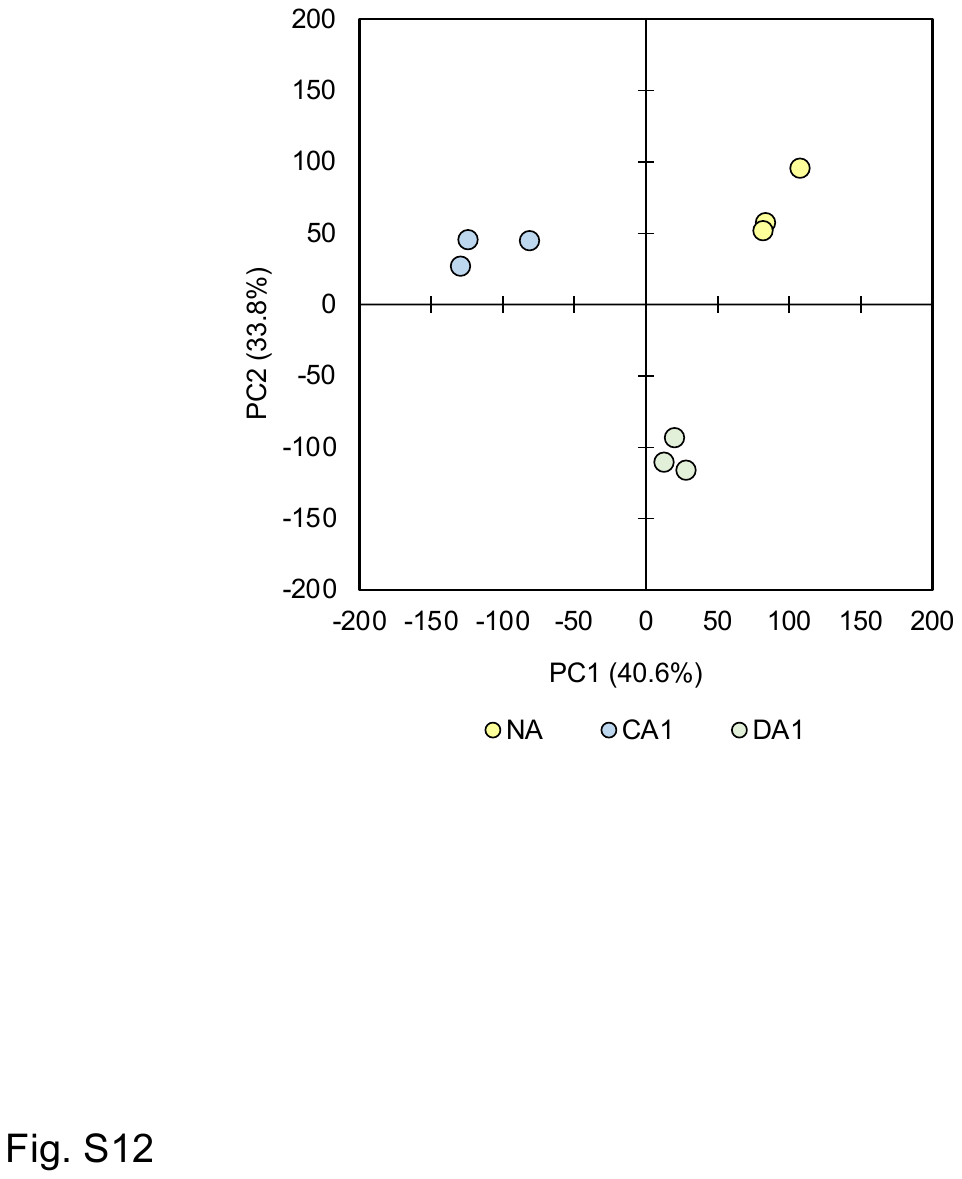

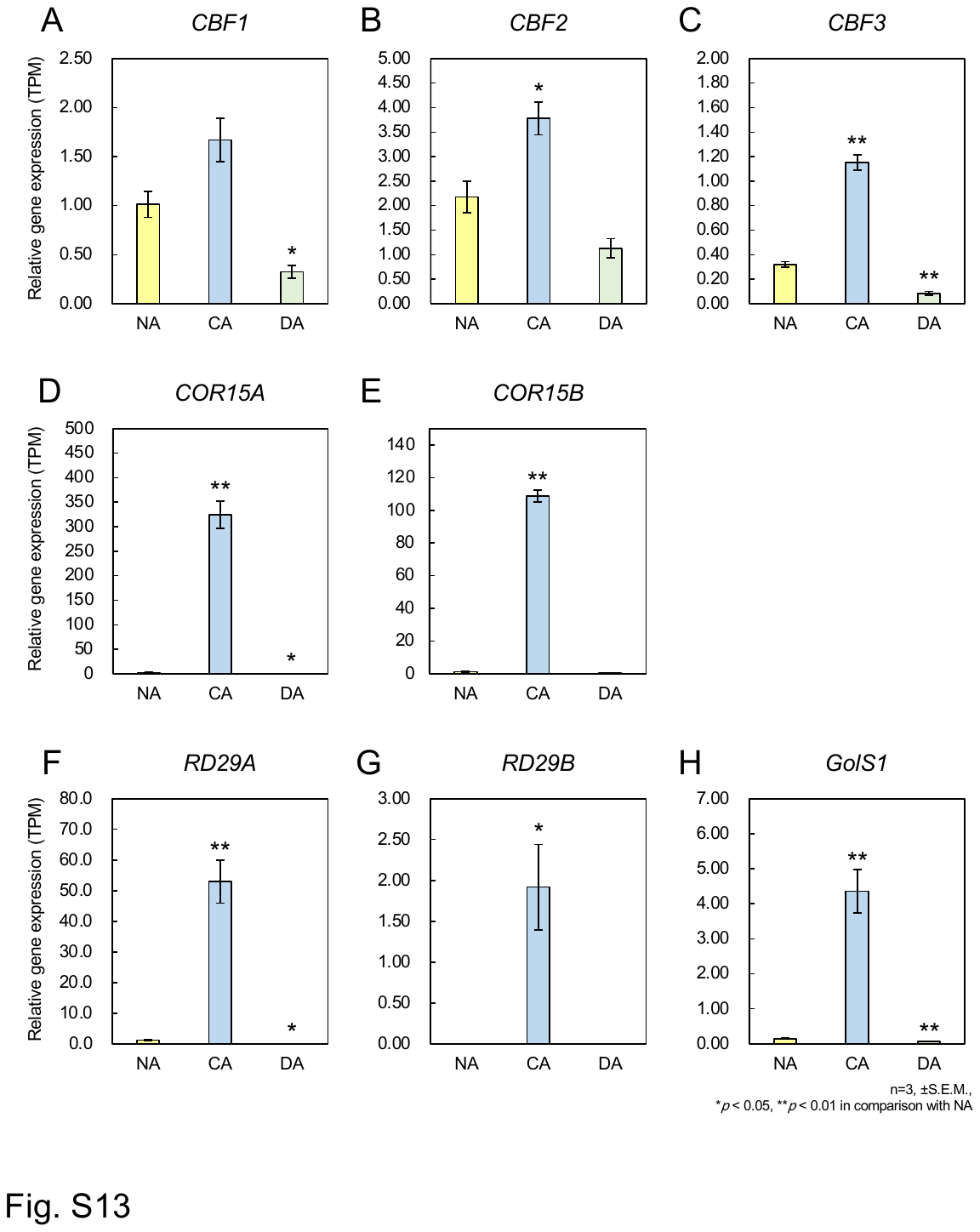
Fig. S13. Expression changes of cold-responsive genes in *Arabidopsis thaliana* during cold acclimation (CA) and de-acclimation (DA). mRNA expression data of all three members of *CBFs* (A-C), *CORs* (D-F), *GolSs* (G and H), and *RD29s* (I and J) were retrieved from RNAseq data (*SI Appendix*, Dataset S1). Relative expression values represent Transcripts Per Million (TPM). Error bars indicate ± S.E.M. (n = 3). Significant differences (Dunnett’s test) between non-acclimation (NA) and other treatment samples are indicated by asterisks above the bars (**p* < 0.05, ***p* < 0.01).

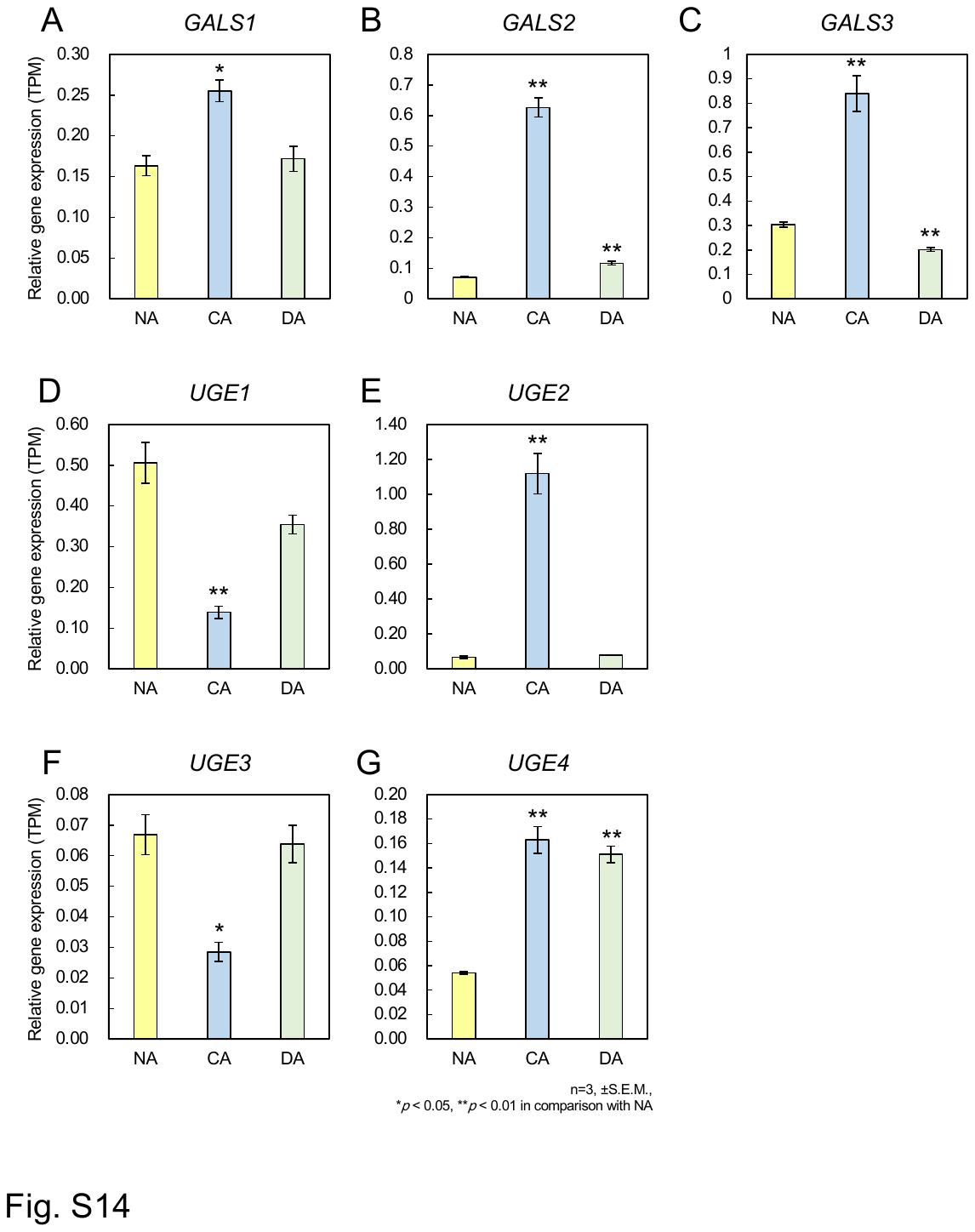
Fig. S14. Expression changes of galactan synthesis-related genes in *Arabidopsis thaliana* during cold acclimation (CA) and de-acclimation (DA). mRNA expression data of all three members of *GALSs* (A-C) and four members of *UGEs* (D-G) were retrieved from RNAseq data (*SI Appendix*, Dataset S1). Relative expression values represent Transcripts Per Million (TPM). Error bars indicate ± S.E.M. (n = 3). Significant differences (Dunnett’s test) between non-acclimation (NA) and other treatment samples are indicated by asterisks above the bars (**p* < 0.05, ***p* < 0.01).

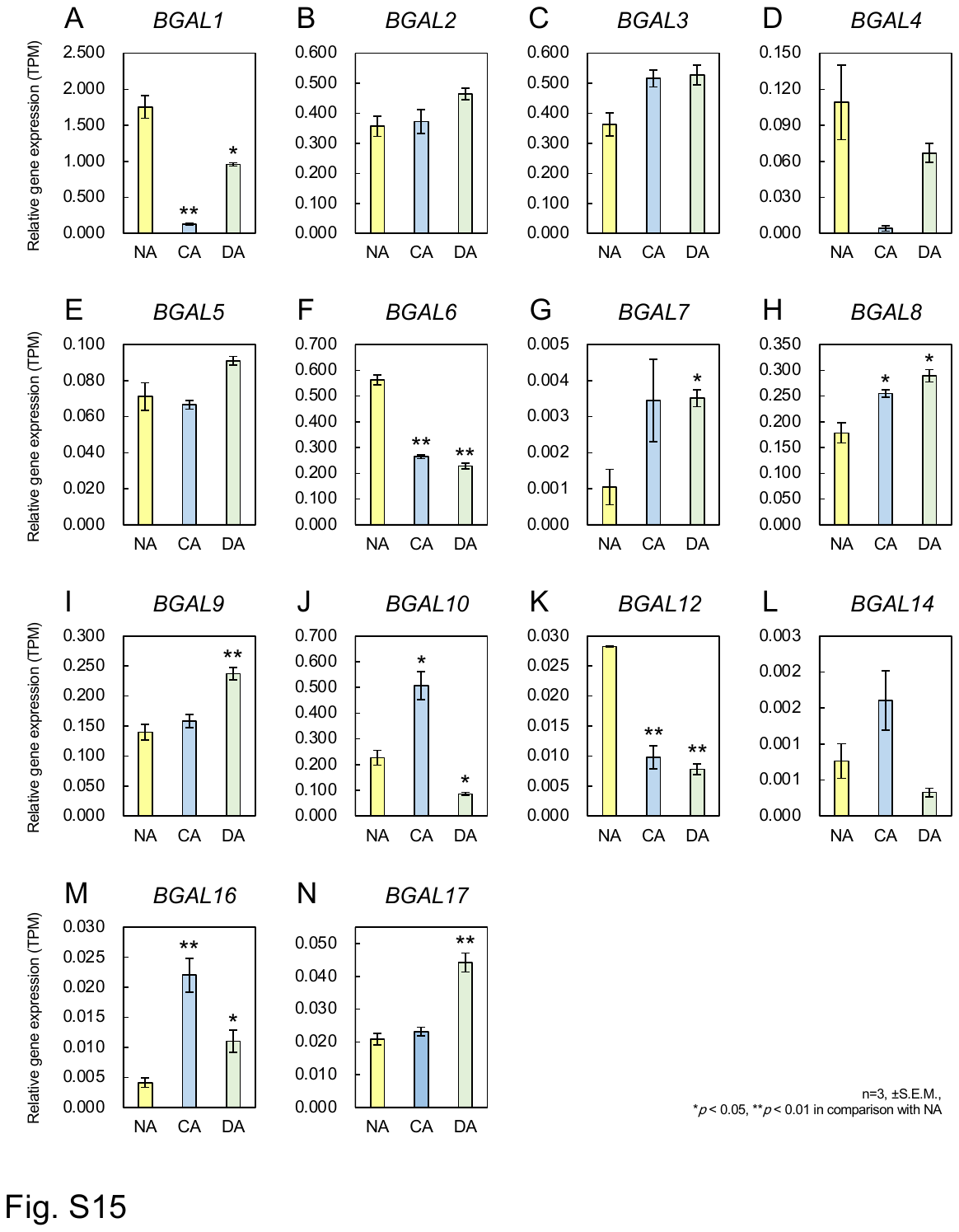
Fig. S15. Expression changes of *β-Galactosidase* (*BGAL*) genes in *Arabidopsis thaliana* during cold acclimation (CA) and de-acclimation (DA). mRNA expression data of 14 members of *BGALs* (A-N) were retrieved from RNAseq data (*SI Appendix*, Dataset S1). Relative expression values represent Transcripts Per Million (TPM). Error bars indicate ± S.E.M. (n = 3). Significant differences (Dunnett’s test) between non-acclimation (NA) and other treatment samples are indicated by asterisks above the bars (**p* < 0.05, ***p* < 0.01).

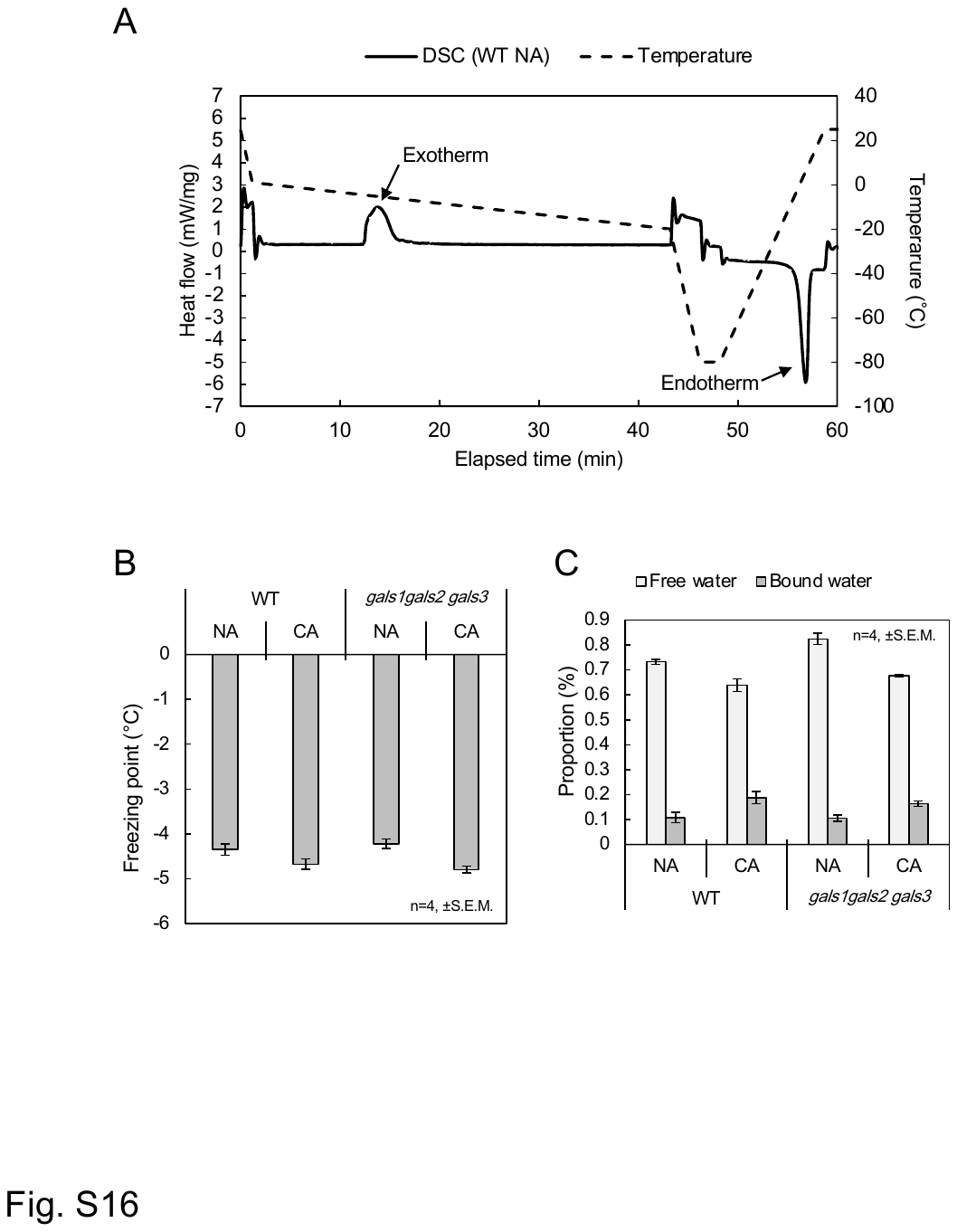
**Fig. S16.** Freezing behavior of leaves detached from wild type (WT) and *gals1 gals2 gals3* analyzed by differential scanning calorimetry (DSC). Typical DSC curve obtained were shown in A. Exotherm due to freezing and endotherm due to thawing of tissue water were observed. The Freezing point (the temperature at which exotherm began to be detected) in each sample is shown in B. Proportions of free water and intermediate water (C) in total water content were calculated from the peak areas of the DSC curves, moisture content and dry weight of the samples used in this study. Error bars indicate ± S.E.M. (n = 4). No significant differences (Student’s *t*-test) between WT and *gals1 gals2 gals3* were found in these results.

Table S1 (separate file). Oligomers used in this study.

Table S2 (separate file). List of sequences used for phylogenetic tree analysis of *GALSs* and *UGEs*

Dataset S1 (separate file). Genes responsive to cold acclimation (CA) and deacclimation (DA) treatments. Genes that changed more than 2-fold or less than 0.5-fold relative to non-acclimated (NA) and less than *p*<0.05 by Dunnett's test for each comparison were listed.
